# Inhibition of IRF4 in dendritic cells by PRR-independent and -dependent signals inhibit Th2 and promote Th17 responses

**DOI:** 10.1101/685008

**Authors:** Jihyung Lee, Junyan Zhang, Young-Jun Chung, Jun Hwan Kim, Chae Min Kook, José M. González-Navajas, David S. Herdman, Bernd Nürnberg, Paul A. Insel, Maripat Corr, Ailin Tao, Kei Yasuda, Ian R. Rifkin, David Broide, Roger Sciammas, Nicholas J.G. Webster, Eyal Raz

## Abstract

Cyclic AMP (cAMP) is involved in multiple biological processes. However, little is known about its role in shaping immunity. Here we show that cAMP-PKA-CREB signaling (a pattern recognition receptor [PRR]-independent) regulates conventional type-2 Dendritic Cells (cDC2s), but not cDC1s and reprograms their Th17-inducing properties via repression of IRF4 and KLF4, transcription factors (TFs) for Th2 induction. Genetic loss of IRF4 phenocopies the effects of cAMP signaling on Th17-induction, indicating that the cAMP effect is secondary to repression of IRF4. Moreover, signaling in cDC2s by a PRR-dependent microbial product, curdlan, represses IRF4 and KLF4, resulting in a pro-Th17 phenotype. These results define a novel signaling pathway by which cDC2s display plasticity and provide a new molecular basis for the novel cDC2 and cDC17 classification. In addition, the data reveal that cAMP signaling can alter DCs function and fate by repressing IRF4 and KLF4, a pathway that can be harnessed for immuno-regulation.

## Introduction

Pattern recognition receptors (PRRs) are germline-encoded proteins expressed primarily on innate immune cells that recognize conserved microbe- or pathogen-associated molecular patterns (PAMPs), and damage-associated molecular pattern (DAMPs). The main paradigm of immune activation posits that triggering of PRRs results in the maturation of DCs and their subsequent activation-acquired properties to elicit and shape CD8^+^ T cells and CD4^+^ Th cells responses (Hansen, Vojtech, & Laing, 2011; Palm & Medzhitov, 2009; Takeuchi & Akira, 2010). DCs, the main antigen presenting cells (APCs), display migratory and functional heterogeneity, and are distributed throughout the body (Guilliams et al., 2014). Conventional DCs (cDCs) are found in most tissues and thought to be of two major lineages, each of which expresses a distinctive set of surface receptors and a unique set of TFs that are necessary for DCs development and function (Guilliams et al., 2014; Murphy, 2013). For example, the splenic cDC1s subpopulation, which promotes CD4^+^ Th1 and CD8^+^ cytotoxic T cells (CTL) responses, requires the TFs interferon regulatory factor 8 (IRF8) and basic leucine-zipper ATF-like transcription factor 3 (BATF3), whereas splenic cDC2s, which promote Th2 and Th17 responses, require IRF4 (Gao et al., 2013; Hambleton et al., 2011; Vander Lugt et al., 2014; Williams et al., 2013) and Kruppel-like factor 4 (KLF4) (Tussiwand et al., 2015). Recent studies have further divided the cDC2s lineage into Th2-inducing (IRF4^+^/KLF4^+^), and Th17-inducing (IRF4^+^/NOTCH2^+^) subpopulations (Bedoui & Heath, 2015; Tussiwand et al., 2015). Although these cDCs express many PRRs to trigger DC maturation, they can elicit and shape the CD4^+^ Th response in the absence of such stimulation. DCs also express multiple G protein-coupled receptors (GPCRs) but their role, in particular of GPCRs that regulate cAMP formation, in DC-related innate immunity and hence, on adaptive immunity, is poorly understood (Idzko, Ferrari, & Eltzschig, 2014; Pearce & Everts, 2015). Thus, in spite of its widely recognized roles in many biological processes, the impact of cAMP on immune responses is not well defined.

IRF4 plays major roles in B and T lymphocyte antigen-dependent responses by controlling the effector properties of expanded clones (De Silva, Simonetti, Heise, & Klein, 2012; Huber & Lohoff, 2014). Induced as an immediate early gene by the antigen receptors, IRF4’s expression scales with the intensity of receptor signaling, thus linking the quality of antigen receptor signaling with B and T cell fate output (Krishnamoorthy et al., 2017; Man et al., 2013; Matsuyama et al., 1995; Nayar et al., 2014; Ochiai et al., 2013; Sciammas et al., 2006; Yao et al., 2013). Differing concentrations of IRF4 induced by antigen receptor signaling are thought to promote differential assembly of IRF4 into distinct TFs and DNA recognition complexes to regulate gene expression programs important for B and T cell fate (Krishnamoorthy et al., 2017; Ochiai et al., 2013). In contrast to other members of the IRF family, expression of IRF4 is not induced by either Type I or Type II interferon (Matsuyama et al., 1995) but rather in response to different modes of NF-kB signaling induced by the antigen receptors, TLR, and TNF receptor systems (Shaffer, Emre, Romesser, & Staudt, 2009). IRF4 plays important roles in controlling the stimulatory properties of DCs; however, the mechanisms and contexts whereby these are deployed and regulate Th responses are less well understood (Bajana, Roach, Turner, Paul, & Kovats, 2012; Vander Lugt et al., 2014).

As we already identified that low levels of cAMP in cDCs promotes Th2 differentiation (Lee et al., 2015) and high levels promotes Th17 (Datta et al., 2010) induction. In this study, we set out to dissect the molecular mechanisms by which cAMP levels regulates these Th responses by cDC1 and cDC2 subsets. We found that increased levels of cAMP in cDC2s, but not cDC1, reprograms a pro-Th17 phenotype (i.e., cDC17s), a process that resulted from a profound inhibition of *Irf4* and *Klf4* expression. We identified that genetic loss of IRF4 in cDC2 accounts for many of cAMP-mediated transcriptional effects that promote pro-Th17 phenotype. Interestingly, the downstream signaling events of this pathway are shared by the activation of a PRR, i.e., Dectin-1, by its microbial ligand, curdlan, which also suppresses the levels of *Irf4* and *Klf4* (Bedoui & Heath, 2015; Tussiwand et al., 2015) and provokes a cDC17 phenotype. Overall, these findings implicate a previously unappreciated DC plasticity provoked by PRR-independent (cAMP) and PRR-dependent (curdlan) signaling that affects Th bias.

## Results

### Cyclic AMP signaling reprograms cDC2 cells from a pro-Th2 to a pro-Th17 phenotype

We previously reported that cAMP signaling in CD11c^+^ bone marrow derived DCs (BMDCs), i.e., BM-derived antigen presenting cells (BM-APCs), affects the differentiation of CD4^+^ T cells (Datta et al., 2010; Lee et al., 2015). Low cAMP levels, as occurs in *Gnas*^ΔCD11c^ mice (generated by breeding of floxed *Gnas* with CD11c-Cre deleter strains), provoke a Th2 polarization that leads to an allergic phenotype (Lee et al., 2015), while cholera toxin (CT) and other treatments that increase cAMP levels in CD11c^+^ cells, induce differentiation to Th17 cells (Datta et al., 2010). Given these observations, it was important to investigate how this phenotypic reprogramming occurs in *bona fide* DCs.

Splenic DCs have been divided into three subsets: two cDCs subsets (cDC1s and cDC2s), and plasmacytoid DCs (pDCs) (Guilliams et al., 2014; Sichien, Lambrecht, Guilliams, & Scott, 2017). We tested the effect of the cell-permeable cAMP analog 8-(4-Chlorophenylthio) adenosine 3’,5’-cyclic monophosphate (CPT) on cDC2s and cDC1s. cDC2s (CD11c^+^CD11b^+^CD8α^-^ splenocytes) (Hey & O’Neill, 2012) were isolated by FACS sorting, pulsed with MHC class II (MHCII) OVA peptide and co-cultured with naïve OVA-specific splenic OT2 CD^4+^ T cells (OT2 cells). Treatment of WT cDC2s with CPT decreased IL-4 (**Figure 1A**) and increased IL-17A concentrations (**Figure 1B**) in co-cultured OT2 cells. IFN-γ and IL-10 concentrations were not changed by CPT-treated cDC2s (**Figure 1C, D**). Analysis of the T cell lineage commitment factors of OT2 cells co-cultured with CPT-treated cDC2s revealed a decrease in *Gata3* and increase in *RORgt* levels (**Figure 1E**). Induction of IL-17 by CPT-treated cDC2 was 2-fold greater in IL-17GFP OT2 cells (**Figure 1F**). The TFs *Irf4* (Gao et al., 2013; Williams et al., 2013) and *Klf4* (Tussiwand et al., 2015) regulate the pro-Th2 phenotype of DCs. Incubation with CPT decreased the expression of *Irf4* and *Klf4* in cDC2s in a time-dependent manner, *Irf8* expression was not affected. To verify that cAMP signaling was activated, we assessed expression of the cAMP-induced gene *Crem* and observed a time-dependent, 10-fold induction (**Figure 1G**). Decreased expression of IRF4 by 50% after CPT treatment was confirmed by intracellular FACS staining (**Figure 1H**).

**Figure 1.**
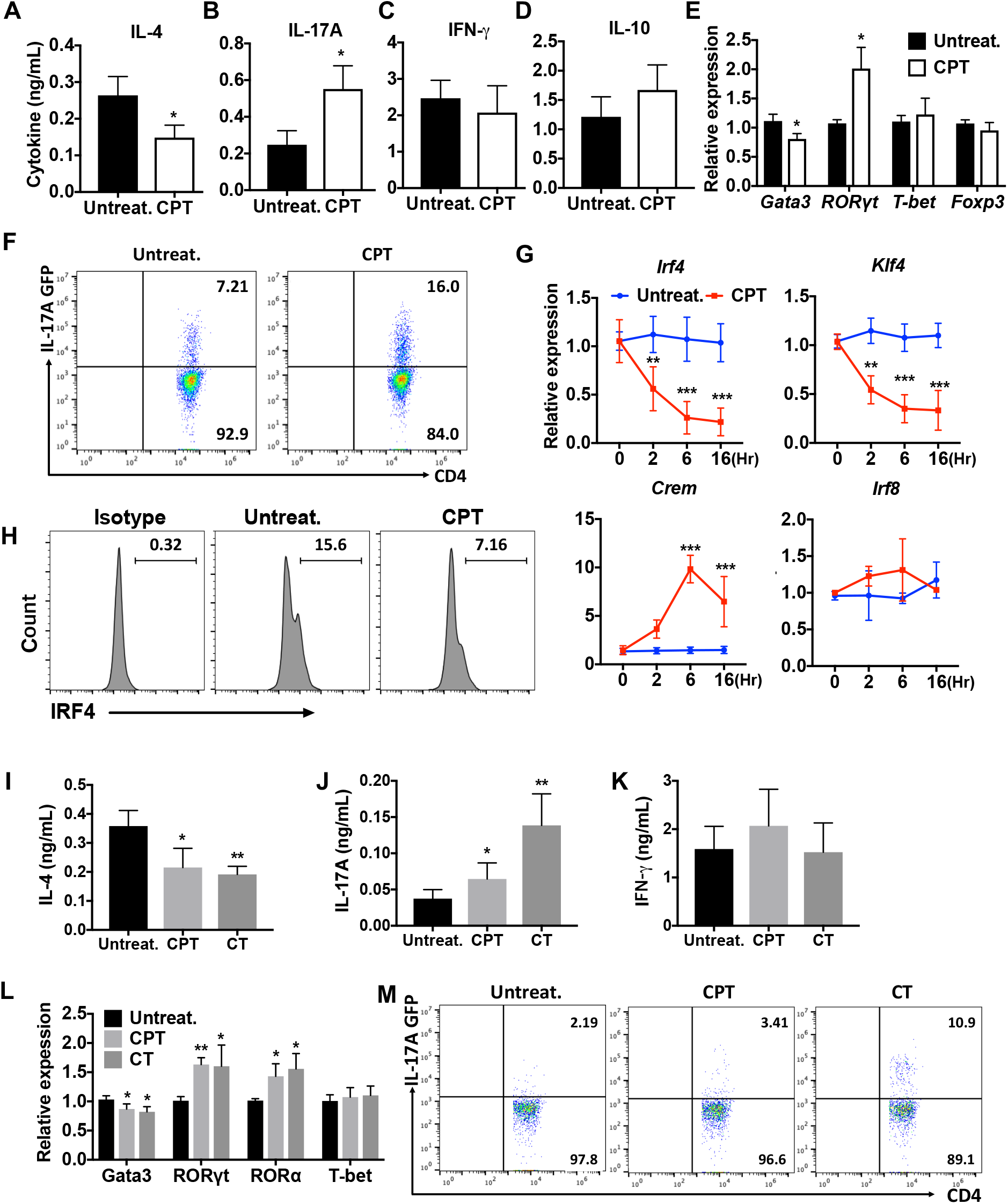
cAMP signaling switches cDC2s to a pro-Th17 bias. **(A-D)** IL-4, IL-17A, IFN-γ and IL-10 levels from anti-CD3/28 Ab-stimulated OT2 cells co-cultured with WT splenic cDC2s (CD11c^+^CD11b^+^CD8α^-^) pretreated with or without CPT. **(E)** QPCR analysis of lineage commitment factors in OT2 T cells co-cultured with WT cDC2s in the presence of CPT. **(F)** GFP expression from IL-17GFP OT2 CD4^+^ T cells co-cultured with WT cDC2s pretreated with or without CPT. **(G)** QPCR of TFs in WT cDC2s treated with CPT (50 µM). Two-way ANOVA with Sidak’s multiple comparisons test; *n*=3 in each group, ** *p*<0.01, \*\*\**p*<0.001. Effect of CPT treatment; *Irf4* (*p*=0.002), *Klf4* (*p*<0.001) and *Crem* (*p*<0.001). **(H)** Intracellular staining of IRF4 in WT cDC2s treated with or without CPT for 48 hr. **(I-M)** Fate mapping: IL-17GFP OT2 CD4^+^ T cells were co-cultured with *Gnas*^ΔCD11c^ BM-APCs to generate memory Th2 cells (1^st^ co-culture). From the 1^st^ co-culture, T1/ST2^+^ cells were FACS sorted and then used for co-culture with WT cDC2s pretreated with or without CPT or Cholera toxin (CT) (2^nd^ co-culture). **(I)** IL-4, **(J)** IL-17A and **(K)** IFN-γ levels, **(L)** QPCR of lineage commitment factors, and **(M)** GFP signal for IL-17 expression in the re-stimulated CD4^+^ T cells from 2^nd^ co-culture. Data are mean ± s.e.m, *n*=3 in each group; * *p*<0.05, ** *p*<0.01, \*\*\**p*<0.001.

Memory Th cells are divided into two subsets: effector memory T (TEM, CD44^+^CD62L^low^) and central memory T (TCM, CD44^+^CD62L^high^) cells (Nakayama et al., 2017). To assess a possible switch in fate from Th2EM to Th17 cells) (Hirota et al., 2011; Nakayama et al., 2017), we first generated Th2EM cells by co-culturing naïve IL-17GFP OT2 cells with *Gnas*^ΔCD11c^ BM-APCs (Lee et al., 2015) and then sorted T1/ST2^+^ cells (Lohning et al., 1998). More than 96% of T1/ST2^+^ cells were Th2EM (data not shown). The sorted cells were then used for a second co-culture with CPT- or cholera toxin (CT) (Datta et al., 2010)-treated WT cDC2s. Th2EM co-cultured with CPT- or CT-treated WT cDC2s had decreased IL-4 (**Figure 1I**), increased IL-17A expression (**Figure 1J**) but unchanged expression of IFN-γ (**Figure 1K**). Th2EM co-cultured with CPT- or CT-treated cDC2s had altered expression of T cell lineage commitment factors: decreased expression of *Gata3* and increased expression of *RORγt* and *RORα* (**Figure 1L**). CPT- or CT-induced Th17 differentiation was confirmed by GFP expression (FACS) in re-stimulated OT2 cells (**Figure 1M**). Expression of T1/ST2 and memory T cell markers, CD62L and CD44, was not changed by the second co-culture (data not shown). Overall, these findings suggest that these cells are akin to Th2-Th17 hybrid cells (Y. H. Wang et al., 2010).

To investigate the possible effect of CPT on the transcriptional program of cDC1s, we isolated CD11c^+^CD11b^-^CD8α^+^ splenocytes and co-cultured them with OT2 cells. Unlike cDC2s, CPT treatment of cDC1s did not change the T cell cytokines and T cell lineage commitment factors in the co-cultured OT2 cells (**Supp. Figure 1A-D**). Furthermore, expression of *Irf4* and *Klf4* was not changed by CPT treatment (**Supp. Figure 1E**) but *Cre*m was induced (**Supp. Figure 1E**). cDC2 cells had 18-fold higher IRF4 (**Supp. Figure 1F**), 3.6-fold higher IRF5 (**Supp. Figure 1G**) and 4.8-fold lower IRF8 (**Supp. Figure 1H**) compared to cDC1s. Expression of IRF5 and IRF8 was not changed by CPT treatment (**Supp. Figure 1I, J**).

To confirm these effects in a genetic model, we isolated CD11c^+^CD11b^+^CD8α^-^ splenocytes from *fl/fl* and *Gnas*^ΔCD11c^ mice (with a prominent decrease in expression of the Gαs protein and in cAMP synthesis and as a result, a bias toward Th2 induction) (Lee et al., 2015) and tested the cDC2 in co-culture with OT2 cells. We found that cDC2s from *Gnas*^ΔCD11c^ mice elicited a 9.7-fold greater IL-4 response, which was completely suppressed by CPT treatment (**Supp. Figure 2A**). IL-17A response induced by CPT-treated *Gnas*^ΔCD11c^ cDC2s increased by 3.4-fold and 2.6-fold in CPT-treated *Gnas*^fl/fl^ cDC2 (**Supp. Figure 2B**). Treatment with CPT suppressed *Gata3* and induced higher *RORγt* levels in co-cultured OT2 cells without altering *T-bet* or *Foxp3* (**Supp. Figure 2D**). Basal *Irf4* and *Klf4* levels in cDC2s were elevated in the *Gnas*^ΔCD11c^ as compared to *Gnas*^fl/fl^ cDC2 but were suppressed in response to CPT (**Supp. Figure 2E, F**). *Irf8* was unchanged and *Crem* was induced by CPT treatment (**Supp. Figure 2E**).

**Figure 2.**
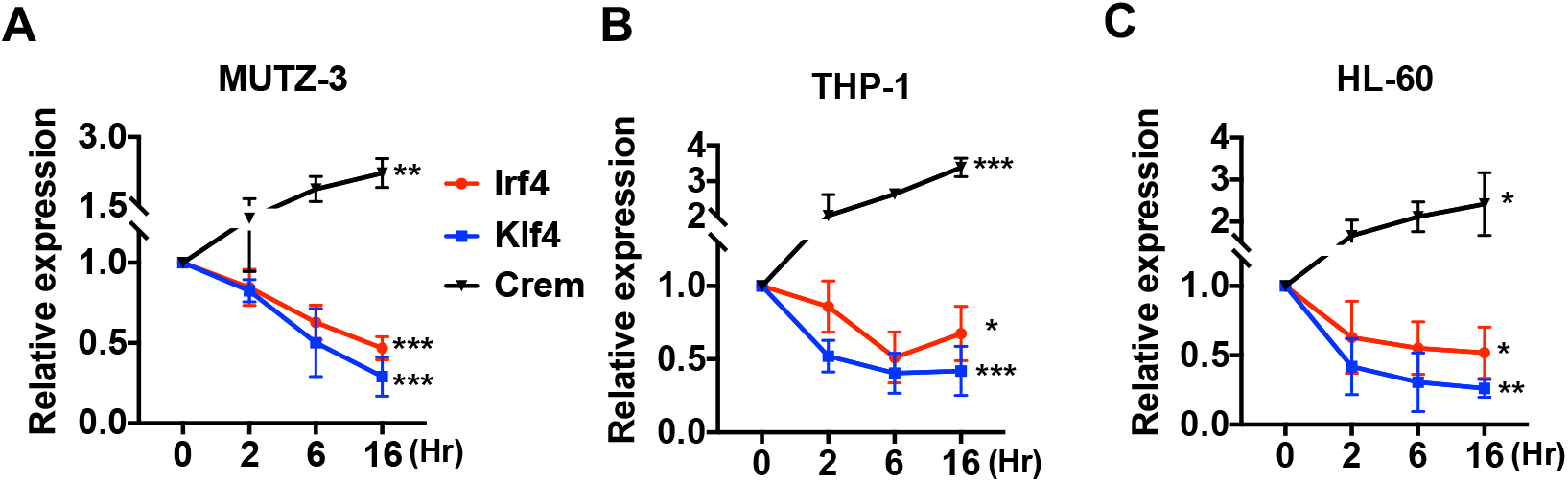
Decreased expression of *Irf4* by CPT in human DC-like cells. DC-like cells were differentiated from **(A)** MUTZ-3, **(B)** THP-1, and **(C)** HL-60 cell lines and treated with CPT for the indicated time. Relative expression of *Irf4*, *Klf4*, and *Crem* were analyzed. Data are mean ± s.e.m, *n*=3 in each group; One-way ANOVA * *p*<0.05, ** *p*<0.01, \*\*\**p*<0.001.

We also checked the effect of cAMP in human DC-like cell lines generated from MUTZ-3 (myelomonocyte) (Mitra et al., 2013), THP-1 (monocyte) (Berges et al., 2005), and HL-60 (promyeloblast) (Koski et al., 1999). As observed with mouse cDC2s or BM-APCs, the human DC-like cells responded to CPT with a decrease in expression of *Irf4* and *Klf4* (**Figure 2**).

To confirm that BM-APCs are a valid model for DC reprogramming and to gain further insight into mechanisms underlying this process, we tested WT CD11c^+^CD135^+^ BM-APCs. We found that different agonists that increase cAMP induced a pro-Th17 phenotype in BM-APCs (**Supp. Figure 3A, B**), a response associated with a 2-fold increase in *RORγt* levels (**Supp. Figure 3C**). *Irf4* and *Klf4* expression were also decreased by cAMP agonist treatment of WT BM-APCs (**Supp. Figure 3D, E**). We also assessed the impact of three cAMP signaling effectors, PKA (Glass, Cheng, Mende-Mueller, Reed, & Walsh, 1989), EPAC (Parnell, Palmer, & Yarwood, 2015), and CREB (Xie et al., 2015), in the inhibition of *Irf4* by PGE2, a Gαs-coupled GPCR agonist (Wehbi & Tasken, 2016). Treatment with Rp-cAMPS, a PKA inhibitor, or 666-15, a CREB inhibitor, but not with CE3F4, an EPAC inhibitor, abolished the PGE2-dependent reduction in *Irf4* expression (**Supp. Figure 3F, G**).

**Figure 3.**
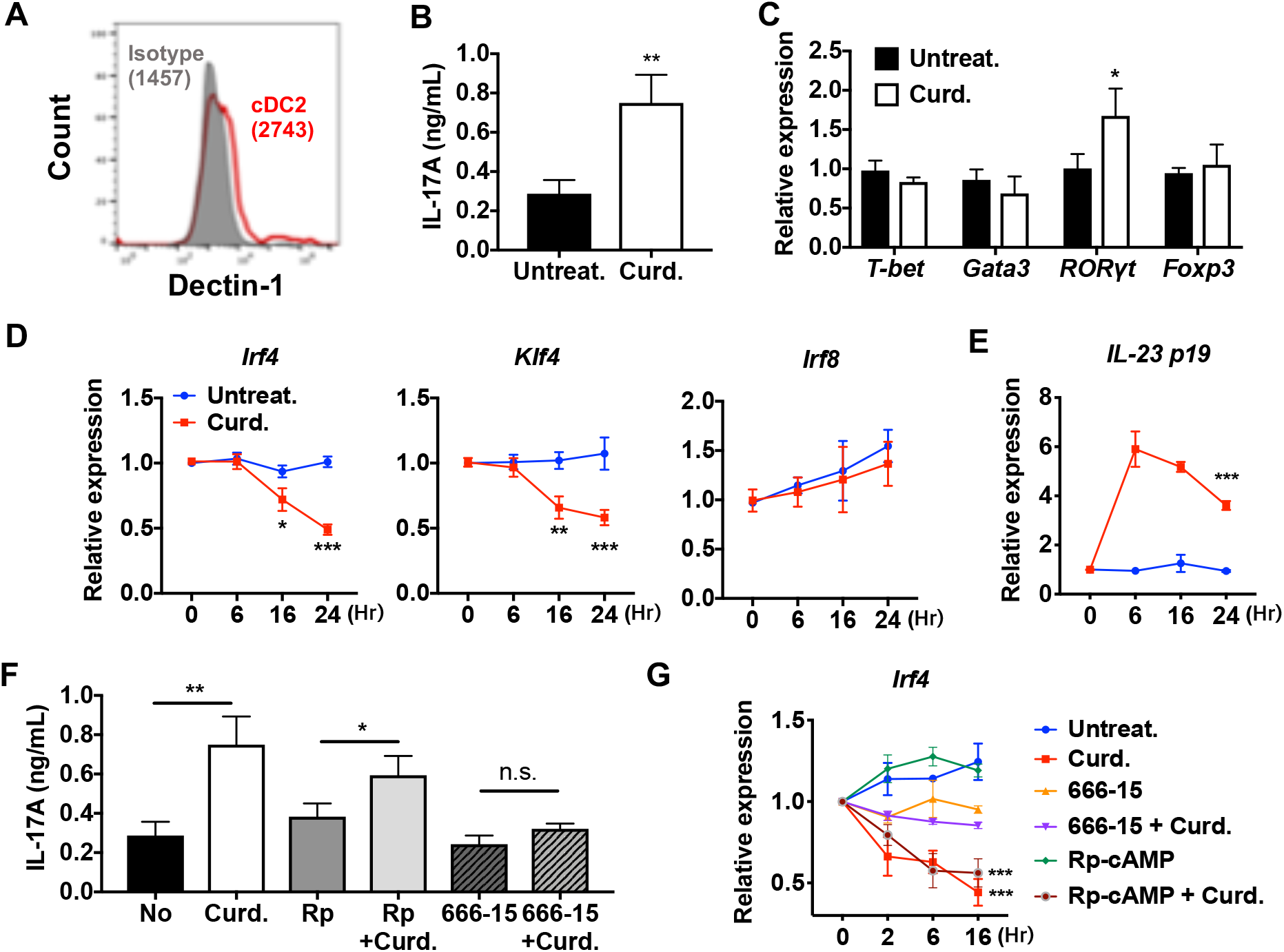
Stimulation of DCs via Dectin-1 (a PPR) regulates IRF4 and KLF4 expression, and induce Th17 differentiation. **(A)** Expression of Dectin-1 on WT cDC2s. Numbers indicate mean fluorescence intensity (GeoMFI). **(B)** IL-17A levels produced by OT2 cells co-cultured with WT splenic cDC2s treated with or without curdlan (10 μg/ml). **(C)** QPCR of lineage commitment factors in OT2 cells co-cultured with curdlan-treated and untreated WT cDC2s. QPCR of **(D)** TFs and **(E)** IL-23 in the WT cDC2s treated with curdlan for the indicated time points. Two-way ANOVA; *n*=3 in each group. **(F)** IL-17A levels produced by OT2 cells co-cultured with WT BM-APC pre-treated with Rp-cAMP (50 µM), or 666-15 (1 µM) 16 hr prior to curdlan treatment. **(G)** QPCR of *Irf4* in WT cDC2s treated with or without curdlan in the presence of inhibitors of CREB (666-15, 1 µM) or PKA (Rp-cAMP, 50 µM). Data are mean ± s.e.m, *n*=3 in each group; * *p*<0.05, ** *p*<0.01, \*\*\**p*<0.001.

Collectively, these results indicate that increased cAMP levels reprogram transcription of cDC2s and BM-APCs from WT and *Gnas*^ΔCD11c^ mice. Low cAMP levels promote expression of a pro-Th2 phenotype while high cAMP levels promote pro-Th17 inducing properties and a subsequent switch from Th2 to Th17 bias. Furthermore, high cAMP levels can override an existing pro-Th2 phenotype and reprogram cDC2 into pro-Th17-inducing cDC17s. This phenotypic switch is mediated by cAMP signaling via PKA-CREB, but not EPAC, and is associated with the inhibition of *Irf4* and *Klf4* in cDC2s.

### PRR-dependent signaling down-regulates IRF4 and KLF4 expression and induces a pro-Th17 phenotype of DCs

To investigate if the reprogramming of WT cDC2 is specific for cAMP signaling, we tested the effect of a microbial Th17 inducer, curdlan. Curdlan is a high molecular weight linear polymer of β (1,3)-glucan derived from soil bacteria and the ligand of the PRR Dectin-1(*Clec7α*) (Yoshitomi et al., 2005). Signaling by Dectin-1 is independent of cAMP and involves recruitment of Syk to the Dectin-1 intracellular tail, followed by activation of MAPK, NF-κB and NFAT (Gross et al., 2006) and induction of IL-23, which promotes Th17 differentiation (LeibundGut-Landmann et al., 2007; Rogers et al., 2005). WT cDC2s express Dectin-1 (**Figure 3A**) and curdlan-pulsed WT cDC2s provoked a 2.5-fold increase in IL-17A (**Figure 3B**) and a 1.7-fold increase in *RORγt* mRNA levels (**Figure 3C**) in co-cultured OT2 cells. Akin to CPT treatment, curdlan also inhibited *Irf4* and *Klf4*, but not *Irf8* expression in cDC2s (**Figure 3D**) and induced *Il-23 p19* mRNA(Agrawal, Gupta, & Agrawal, 2010) by 6-fold (**Figure 3E**). Moreover, curdlan-treated cDC2s increased IL-17A, a response blocked by the CREB inhibitor, 666-15, but not by the PKA inhibitor, Rp-cAMP (**Figure 3F, G**). Unlike the response of cDC2s, cDC1s cells did not express, or express very low levels of Dectin-1 (**Supp. Figure 4A**). Indeed, curdlan treatment of cDC1s neither change the T cell cytokines in co-cultured OT2 cells (**Supp. Figure 4B, C**), nor the expression of *Irf4* and *Il-23 p19* mRNA in cDC1s (**Supp. Figure 4D, E**). Overall, these data indicate that curdlan acts via its PRR to induce a Th17 bias by cDC2s via a CREB-dependent, cAMP-PKA-independent pathway. Furthermore, they define a convergence of signaling of these two pathways on CREB that allows for phenotypic plasticity of DCs and provide a new molecular basis for the classification of novel cDC2 and cDC17 subsets.

**Figure 4.**
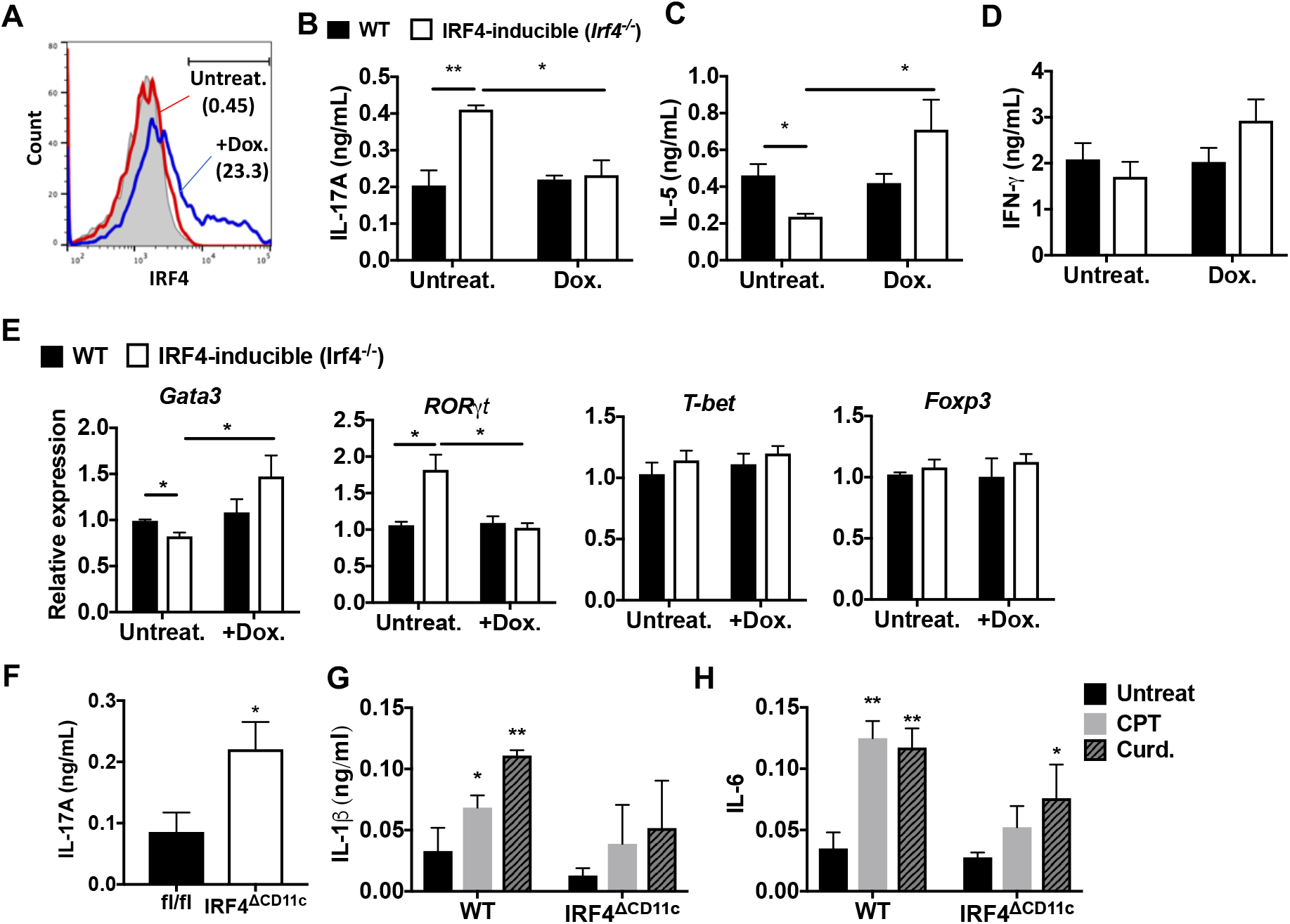
Decreased IRF4 expression in cDC2s promotes pro-Th17 phenotype. **(A)** IRF4 expression in the *Irf4*-*inducible* (*Irf4^-/-^*) cDC2s treated with or without doxycycline (Dox, 200 ng/ml) for 16 hr. **(B-D)** IL-17A, IL-5 and IFN-γ levels and **(E)** T cell lineage commitment factors from the re-stimulated OT2 cells co-cultured with cDC2s from WT and *Irf4*-*inducible* (*Irf4^-/-^*) mice under the conditions described above. **(F)** IL-17A levels from anti-CD3/28 Ab-stimulated OT2 cells co-cultured with IRF4^ΔCD11c^ cDC2s. **(G)** IL-1β and **(H)** IL-6 level in WT and IRF4^Δ^^CD11c^ cDC2s after treatment of CPT or Curdlan for 24 hr. Data are mean ± s.e.m, *n*=3 in each group; * *p*<0.05, ** *p*<0.01.

### Loss of IRF4 or IRF5 in DCs promotes or inhibits, respectively, the pro-Th17 DC phenotype and subsequent Th17 bias

IRF4 is a key TF in the development and function of innate immune cells (macrophages and cDCs) and adaptive immune cells (B and T cells) (Biswas et al., 2012; Mittrucker et al., 1997; Satoh et al., 2010; Suzuki et al., 2004). IRF4-deficient cDCs display dysfunctional antigen processing and presentation (Vander Lugt et al., 2014) and do not migrate to draining lymph nodes, which may contribute to their inability to stimulate Th2 or Th17 responses (Bajana et al., 2012; Vander Lugt et al., 2014). The data above indicate that cAMP signaling in splenic cDC2s inhibits IRF4 expression and provokes a Th17 bias. To determine if inhibition of IRF4 levels affects the pro-Th17 phenotype, we used splenic cDC2s from mice with genetic deletion and conditional re-expression of IRF4. These mice harbor a tetracycline (tet)-inducible cDNA allele of *Irf4* and the M2rtTA tet-activator allele, hereafter termed *Irf4-inducible* (*Irf4^-/-^*) mice. The tetracycline analog Doxycycline (Dox) induces IRF4 expression in cDC2s from these mice (Ochiai et al., 2013) (**Figure 4A**). Remarkably, unlike WT cDC2s, *Irf4-inducible* (*Irf4^-/-^*) cDC2s pulsed with MHCII OVA peptide (i.e., without any additional stimulation) increased IL-17A and decreased IL-5 production in co-cultured OT2 cells (**Figure 4B, C**). In contrast, activation of IRF4 with Dox treatment inhibited IL-17A production (**Figure 4B**) and increasedIL-5 levels (**Figure 4C**); IL-4 was not detected and IFN-γ was not affected by the IRF4 expression in cDC2s (**Figure 4D**). We observed altered expression of the of T cell lineage commitment factors *Gata3* (which was increased) and *RORγt* (which was decreased) in OT2 cells co-cultured with Dox-treated *Irf4-inducible* (*Irf4^-/-^*) (**Figure 4E**). cDC2s from *IRF4*^ΔCD11c^ mice also induced Th17 differentiation spontaneously alone (**Figure 4F**), or upon treatment with CPT or curdlan. However these cells did not produce significant levels of IL-1β or IL-6, i.e., established cytokines that provoke Th17 differentiation (Chen & O’Shea, 2008) (**Figure 4G, H**).

Since IRF5 in macrophages has an inductive role in Th17 response (Krausgruber et al., 2011), we tested whether IRF5 in cDC2s affects the pro-Th17 phenotype. CPT treatment of OVA-pulsed cDC2 from *Irf5 ^-/-^* mice did not provoke a Th17 response (**Supp. Figure 5A**): even though *Irf4* and *Klf4* levels were inhibited, *Irf8* was unchanged and *Crem* was induced (**Supp. Figure 5B**). Thus, even in the absence of IRF4, IRF5 was required for the cAMP-induced pro-Th17 phenotype. *Irf5* mRNA levels were similar in *Irf4^-/-^* and WT mice, indicating that IRF5 expression is independent of IRF4.

**Figure 5.**
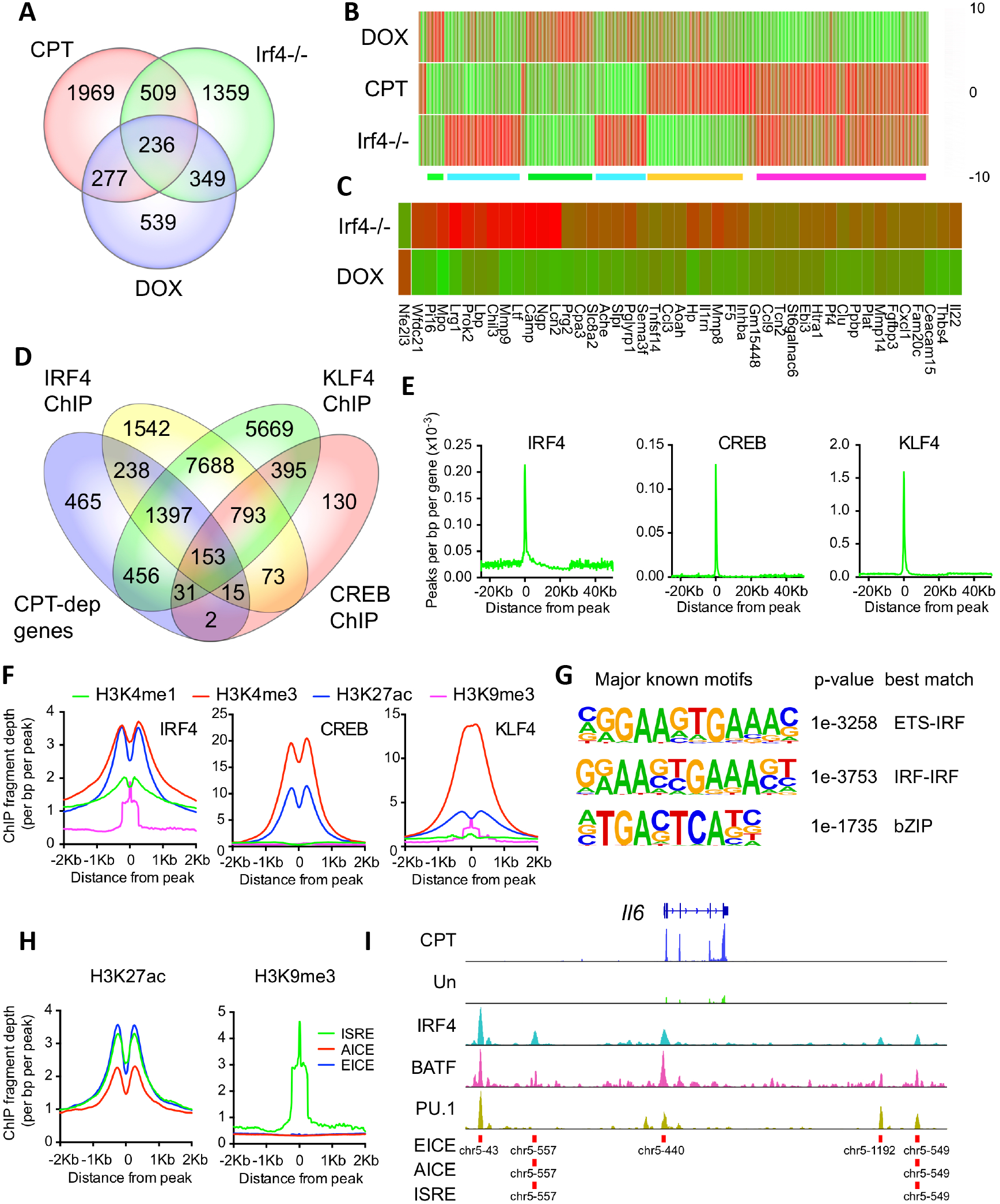
CPT and IRF4 transcriptomic effects and analysis of genome-wide binding of IRF4, CREB and KLF4. **(A)** Venn diagram showing overlap of genes altered by CPT treatment (50 µM, 16 h) of WT splenic cDC2s, with splenic cDC2s derived from *Irf4*^-/-^ mice, and splenic cDC2s from *Irf4^-/-^* mice that have been induced with Dox (200 ng/ml, 16 h). **(B)** Heatmap showing expression (log2 fold-change) of the 745 genes common to CPT and *Irf4*^-/-^ cDC2s. Colored bars under the heatmap indicate clusters of genes with similar expression patterns. **(C)** Heatmap showing expression (log2 fold-change) of secreted genes that are altered in *Irf4^-/-^* and Dox datasets. **(D)** Venn diagram showing overlap of CPT-dependent genes with ChIPseq peaks for IRF4, CREB1 and KLF4. **(E)** Metagene analysis of the localized binding of IRF4, CREB1 and KLF4 to a synthetic gene. The synthetic gene is 25kB in length and is flanked by 25 kB of upstream and downstream sequence. The transcriptional start site is indicated at 0. The plot shows the number of peaks per bp per gene (x10^-3^). **(F)** Co-localization of histone epigenetic modifications H3K4me1, H3K4me3, H3K27ac and H3K9me3 at IRF4, CREB and KLF4 peaks. The graphs show the ChIP fragment depth relative to the center of the TF peak. **(G)** Known transcription factor motifs identified in the IRF4 binding peaks. Height of the letter indicates its conservation. The p-value for the motif and its best match are shown. **(H)** Co-localization of H3K27ac and HeK9me3 modifications at the peaks with the three IRF4 motifs (ISRE, EICE, AICE). **(I)** IRF4, BATF, and PU.1 binding to the IRF4-super enhancer at the *Il6* locus. The *Il6* gene structure is shown at the top and RNAseq reads from untreated and CPT-treated cDC2s are shown in green and blue. Locations of individual motifs are indicated below the binding.

Overall, these results emphasize that the deletion or substantial inhibition of IRF4 levels in DCs is necessary but not sufficient for provoking Th17 differentiation, and that IRF5 expression in DCs is necessary but insufficient for cAMP to provoke Th17 bias.

### IRF4 mediates cAMP-promoted transcriptional events

To assess mechanisms for cAMP-PKA-mediated suppression of IRF4 expression, we initially performed transcriptional profiling (RNAseq) of untreated or CPT-treated splenic cDC2s from WT mice, global *Irf4* KO mice (*Irf4^-/-^*), and *Irf4*-*inducible* (*Irf4^-/-^*) mice treated with Dox. Treatment of splenic cDC2s from WT mice with CPT altered expression of 2991 genes (FDR <0.05, **Figure 5A**), results resembling those previously observed for BM-APCs (Datta et al., 2010). Expression of 2454 genes was significantly altered in *Irf4^-/-^* cDC2s compared to WT cDC2s but only 1401 genes had altered expression in *Irf4^-/-^* cDC2s with restored expression of *Irf4* (Dox) (**Figure 5A; Supp. Tables 1-3**). Surprisingly, only 24% of the genes altered in the *Irf4*^-/-^ cDC2s were restored by re-expression of IRF4 in Dox-treated *Irf4*-*inducible* (*Irf4^-/-^*) cells, and only 42% of the Dox-dependent genes were altered in the *Irf4*^-/-^ cells. These results are not related to the stringency of the multiple testing correction as the discordance remained (30% and 46%, respectively) even without a correction for FDR. This observation is consistent with prior data for genes that display a bimodal pattern of expression as a function of IRF4 concentration in B and T cells (Iwata et al., 2017; Krishnamoorthy et al., 2017; Ochiai et al., 2013).

Transcriptional network analysis showed that the predominant TF driving CPT-dependent genes was CREB1, which appeared to regulate 1134 genes (38%, z-score 236) (**Supp. Table 4**). Unexpectedly, CREB1 was also the major TF “driver” of genes altered in *Irf4*^-/-^ cells and Dox-treated *Irf4*-*inducible* (*Irf4^-/-^*) cells (592 and 442 such genes, 24% and 32%, z-scores 233 and 240, respectively) (**Supp. Table 5 and 6**). These observations implicate CREB1 as a significant cofactor for genes regulated by IRF4 and suggest a link between cAMP signaling and IRF4-regulated transcription in DCs.

The genes altered by CPT treatment or loss of IRF4 significantly overlapped (p<0.0001 by Chi-square): 745 genes (**Supp. Table 7**) were in common (**Figure 5A**), a greater number than between the *Irf4^-/-^* and Dox-treated cells (585 genes, **Supp. Table 8**). To gain insight into the gene expression changes, clustering analysis was performed and revealed a number of distinct classes of changes among the 745 common genes, as shown in a heatmap (**Figure 5B**). A large cluster of genes was concordantly induced by CPT or loss of IRF4, and reduced by IRF4 re-expression (**Figure 5B magenta**); a smaller cluster of genes was reduced by CPT or IRF4 loss, and restored by IRF4 re-expression (**Figure 5B green**). These two clusters may indicate genes indirectly regulated by cAMP via its suppression of IRF4 expression. Examples of genes in these clusters are the IL-1 receptor antagonist (*Il1rn)* and C-C motif chemokine receptor 2 (*Ccr2)* (**Supp. Figure 6A, B**). Other clusters are discordant. i.e., decreased by CPT but increased in the *Irf4^-/-^* cells or vice versa (**Figure 5B, green and orange**). Examples of genes in these clusters are plasminogen activator, urokinase receptor (*Plaur)* and matrix metalloprotease 27 (*Mmp27*) (**Supp. Figure 6C, D**).

**Figure 6.**
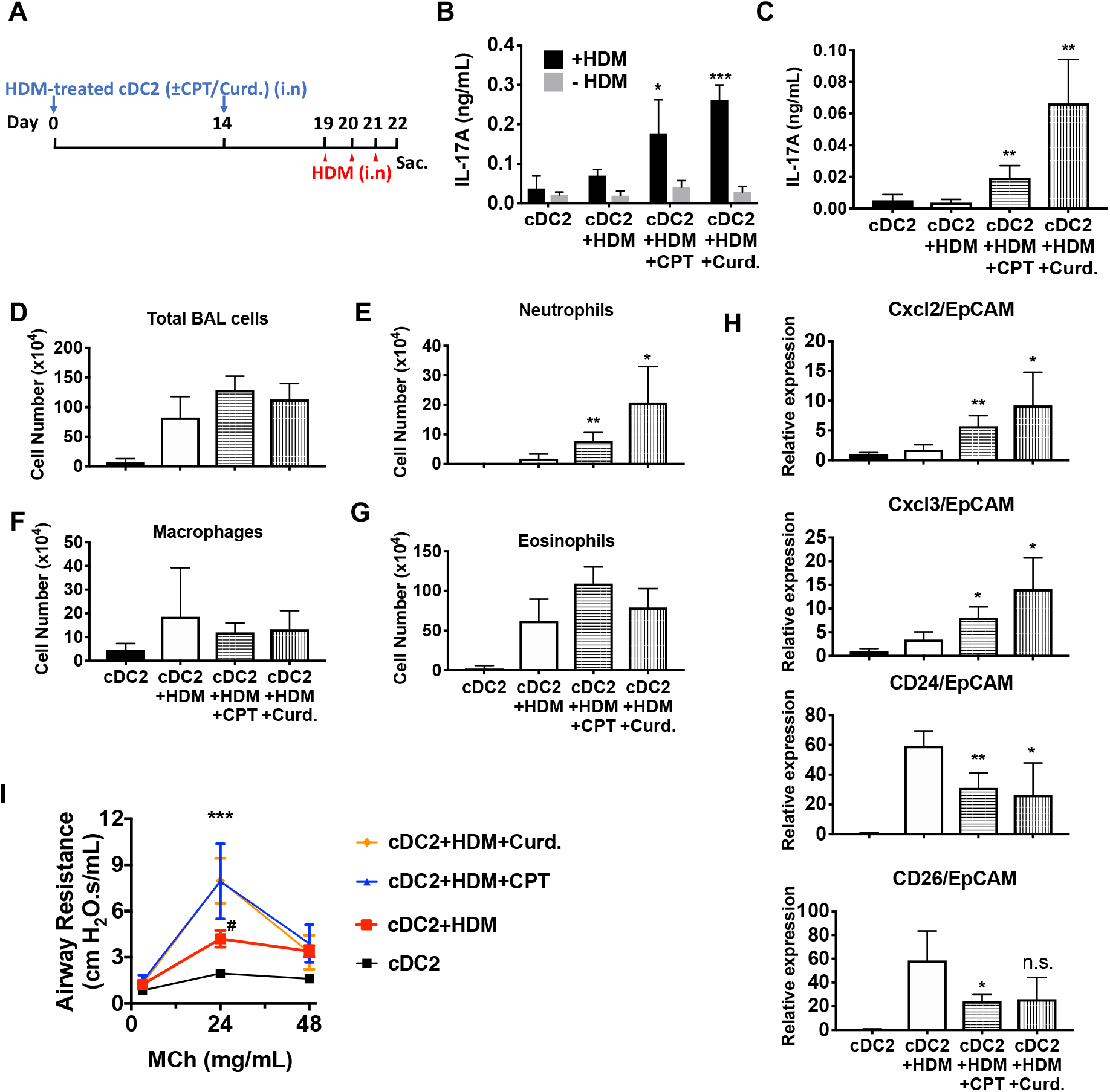
Adoptive transfer of HDM-pulsed, CPT or curdlan-treated cDC2 induces a Th17 bias and neutrophil infiltration in WT recipients. **(A)** Schematic of adoptive transfer protocol. WT cDC2s were incubated with HDM (50 μg/ml) in the presence of CPT or curdlan prior to i.n. transfer to WT mice (B6 mice, 1×10^6^ cells/recipient) on day 0 and 14. HDM (12.5 µg/mouse) was used for the i.n. challenge on days 19, 20 and 21. On day 22, lungs from each group were harvested and processed a single cell suspension. **(B)** IL-17A level in the HDM (50 μg/ml)-stimulated lung cells. BAL fluid was analyzed for **(C)** IL-17 level (ELISA). **(D)** Total cells **(E)** neutrophils, **(F)** macrophages and **(G)** eosinophils number were counted in the BAL fluid. **(H)** Relative expression of Neut (Cxcl2 and Cxcl3)- and / Eos (CD24 and CD26)-infiltration related genes in the lung tissue. Expression of each gene was normalized by expression of epithelial specific housekeeping gene, EpCAM. **(I)** Airway resistance after MCh challenge was measured in the various experimental groups. Data are mean ± s.e.m, *n*=4 recipient mice in each group; * *p*<0.05, ** *p*<0.01, \*\*\**p*<0.001.

Since CPT treatment, loss of IRF4 or restoration by Dox treatment all changed the ability of cDC2s to promote Th bias, we reasoned that key mediators of this bias would be among the 236 genes that were common to the three datasets (**Supp. Table 9**). Transcriptional network analysis and enrichment analysis for pathways and processes of these 236 common genes revealed that the most significantly enriched TF was CREB1 (**Supp. Table 10**) and the most significantly enriched pathways related to cell adhesion, CCL2 signaling, immune cell migration, myeloid differentiation, neutrophil activation, and inter-cellular interactions in COPD (**Supp. Table 11**). Network enrichment of processes centered on angiogenesis and immune cell activation/response pathways (**Supp. Table 11**). The two highest enriched GO processes were secretion-related (p<10^-25^), suggesting involvement of secreted mediators of Th differentiation (**Supp. Table 11**). We therefore intersected this group of genes with curated and highly-likely predicted secreted proteins from the MetazSecKB database (Meinken, Walker, Cooper, & Min, 2015) and found that secreted proteins were highly enriched (59 of the 236 common genes (25%), p<0.0001). We subdivided these genes into those with concordant or discordant regulation between CPT treatment and *Irf4*^-/-^ (**Supp. Table 12**). Of the 22 concordant genes among the data sets, all were up-regulated by treatment with CPT or in *Irf4*^-/-^, suggesting that promotion of Th17 differentiation requires production of a Th17 mediator rather than loss of a Th2 mediator. Since loss of *Irf4* alone can lead to Th17 differentiation, we inspected the secreted genes for those induced by IRF4 loss and restored by IRF4 “rescue” (**Figure 5C**). We also intersected the differentially expressed genes with predicted plasma membrane proteins and found that these are significantly enriched (82 of the 236 common genes (34%), p<0.0001) (**Supp. Table 13**). These results indicate that many of the cAMP-dependent transcriptional changes may be a consequence of the repression of IRF4 expression. Furthermore, the IRF4- and cAMP-dependent genes are enriched in secreted and plasma membrane proteins that could modulate T cell differentiation.

### Transcriptional changes correlate with transcription factor binding but not with alteration in chromatin accessibility

To gain a better understanding of the genomic events involved in these transcriptional changes, we performed ATACseq for changes in chromatin accessibility. No significant differences were observed in ATACseq peaks for any of the sample groups, indicating that the cDC2 plasticity is not accompanied by gross changes in chromatin structure (**Supp. Figure 6**). We then analyzed ChIPseq datasets derived from IRF4, CREB, KLF4, and histone modifications in BMDC, cDC2 and related cells (**Suppl. Table 14**). We examined the co-localization of binding sites for IRF4, CREB and KLF4 within CPT-regulated genes (**Figure 5D**). Surprisingly, only 7% of CPT-regulated genes (n=201) contained CREB binding sites, in contrast to 65% with an IRF4 binding site, 74% with a KLF4 binding site, and 56% with both IRF4 and KLF4 binding sites (**Figure 5D**). Combining the expression data with the ChIP data showed that the presence of neither a CREB nor a KLF4 binding site was predictive of up- or down-regulation of expression by CPT treatment (p=1.0 and 0.82, and p=0.97 and 0.68, respectively) (**Supp. Figure 7A**). By contrast, the presence of an IRF4 binding site was predictive of both up- and down-regulation by CPT (p=3.6×10^-5^ and 2.6×10^-6^, respectively), implying that cAMP regulation of genes in cDC2 may be mediated by alterations in IRF4 expression. Indeed, 890 of the 2991 CPT-regulated genes (30%) were also regulated by IRF4 loss or over-expression (**Figure 5A**). A similar analysis with genes altered in the *Irf4^-/-^* and *Irf4*-*inducible* (*Irf4^-/-^*) cells revealed that IRF4-regulated genes in those cDC2s were not predicted by the presence of a CREB binding site (**Supp. Figure 7B, C**), but that up-regulated genes in the *Irf4^-/-^* cells and down-regulated genes in the *Irf4*-*inducible* (*Irf4^-/-^*) cells were highly predicted by IRF4 binding sites (p=6.9×10^-13^ and p=1.7×10^-14^, respectively). IRF4 binding sites were less predictive for genes that were decreased in *Irf4^-/-^* cells (p=0.0012) but not were highly predictive for genes induced by IRF4 over-expression (p=7×10^-6^). The presence of KLF4 binding sites predicted genes that decreased with IRF4 overexpression (p=7.7×10^-5^).

To further understand the transcriptional regulation, we analyzed the location of the IRF4, CREB and KLF4 binding sites relative to the transcriptional start sites (TSS) and the gene bodies, as well as 25Kb upstream and downstream of the genes. Both CREB and KLF4 primarily bound to the TSS with very little binding to the gene body, upstream or downstream regions. In contrast, IRF4 showed weaker binding to the TSS but more substantial binding upstream and downstream of the genes (**Figure 5E**). Consistent with this observation, the IRF4, CREB and KLF4 binding sites were associated with H3K4me3 and H3K27ac marks of transcriptionally active promoters (Jenuwein & Allis, 2001) but only IRF4 binding sites were associated with the H3K4me1 modification, which is indicative of enhancers, and with the repressive H3K9me3 modification in the middle of the IRF4 peak (**Figure 5F**).

We investigated the known TF motifs bound by IRF4 and found that they fell into three categories: 50% of the sites had ETS:IRF motifs similar to proposed EICEs (Brass, Kehrli, Eisenbeis, Storb, & Singh, 1996; Ochiai et al., 2013), 24% had duplicated IRF motifs similar to ISREs (Bovolenta et al., 1994; Ochiai et al., 2013), and 23% had bZIP motifs similar to AICEs (**Figure 5G and Supp. Figure 8A**). The sub-populations of IRF4 binding sites had similar association with H3K4me1 and H3K27ac enhancer marks but different co-localizations with histone repressive marks, in that only IRF4 at ISRE sites was associated with the transcriptionally repressive H3K9me3 mark (**Figure 5H**). IRF4 may thus regulate gene expression in a context-specific manner as well as by differential binding dynamics dependent on its interactions with its binding partners (Krishnamoorthy et al., 2017; Ochiai et al., 2013).

Given this difference in association with transcriptionally repressive histone modifications, we performed a similar analysis with transcriptional changes in response to treatment with CPT (**Supp. Figure 8B**). The presence of an AICE most strongly predicted up-regulation by CPT treatment (p=1.7×10^-6^), whereas all three response elements predicted up-regulation in *Irf4^-/-^* cells, and down-regulation in Dox-treated *Irf4* cells. Thus, all three sites can mediate repression by IRF4. The AICE and ISRE, but not the EICE, also predicted gene induction in the *Irf4*-*inducible* (*Irf4^-/-^*) cells, suggesting that these two elements can confer gene induction by IRF4 (**Supp. Figure 8B**). Many of the IRF4 sites clustered together. Using the Young lab criteria, we found evidence for 106 super enhancers with multiple IRF4 binding sites (**Supp. Table 15**) (Whyte et al., 2013). Inspection of these super enhancers revealed that one was located in the *Il6* locus and had multiple binding sites for IRF4, BATF and PU.1; these peaks corresponded to EICEs, AICEs and ISREs identified in the motif analysis (**Figure 5I**). Localization of IRF4, CREB, KLF4, H3K4me3 and H3K27ac peaks and the presence of open chromatin (measured by ATACseq) in the vicinity of the *Irf4* and *Klf4* genes indicated that the *Klf4* gene, but not the *Irf4* gene, may be a direct target for CREB (**Supp. Figure 9**).

Thus, these genomic data were consistent with a role for IRF4 binding to and regulation of both cAMP- and IRF4-dependent genes. The results also suggested that CREB did not directly regulate many of these cAMP-dependent genes. The data were consistent with the notion that IRF4-dependent transcriptional events are modulated by the partner TF as IRF4 homodimers or heterodimers with ETS or bZIP proteins predicted gene repression, but only IRF4 homodimers and bZIP heterodimers mediate gene induction.

### Adoptive transfer of HDM-pulsed, CPT- or curdlan-treated cDC2s induce a Th17 bias and neutrophilic infiltration

We hypothesized that activation of DCs via the cAMP pathway would modify their Th-inducing properties *in vivo*. To test this hypothesis, we evaluated the impact of CPT on DCs in the induction of Th17 bias by using a DC-based adoptive transfer (Lambrecht & Hammad, 2012) model (**Figure 6A**). We used HDM- and curdlan-treated cDC2s, as negative and positive controls, respectively, of Th17 induction. Intranasal (i.n.) transfer of HDM-pulsed, CPT-treated or curdlan-treated WT cDC2s followed by HDM i.n challenge increased the production of IL-17A in the lung and airway of WT recipient mice (**Figure 6B, C**). However, we did not observe differences in IL-4 (>0.04 ng/ml) and IFN-γ (>0.02 ng/ml) levels in the various groups (data not shown). We also detected an increased number of neutrophils in the bronchoalveolar lavage (BAL) fluid (**Figure 6D-G**) and expression of CXCL2 and CXCL3, whose expression is intimately related to neutrophilic infiltration in the CPT- or curdlan-treated cDC2 groups (**Figure 6H**). Expression of eosinophilic infiltration-related genes, CD24 and CD26, was decreased in those groups (**Figure 6H**). Moreover, CPT- or curdlan-treated WT cDC2s transferred to WT recipient mice had increased airway resistance (**Figure 6I**). We also checked the effects of CPT or curdlan on OVA-pulsed cDC2s *ex-vivo* prior to their i.n transfer and found a similar pattern to that observed for HDM (**Supp. Figure 10**). Thus, induction of Th17 in the lungs by CPT or curdlan with two different antigen systems yielded a shift from a Th2 toward Th17 pulmonary response with a mixed eosinophil/neutrophil inflammatory infiltrate.

## Discussion

The current paradigm for the activation and maturation of DCs emphasizes the dominant role of PRRs. Such activated DCs are characterized by efficient antigen processing and presentation, up-regulation of co-stimulatory molecules, and secretion of cytokines and other immunomodulatory factors, most of which contribute to the differentiation of naïve T cells to effector subpopulations, such as CTL or Th subsets (Akira, Uematsu, & Takeuchi, 2006; Merad, Sathe, Helft, Miller, & Mortha, 2013). Our results indicate that cAMP via a PRR-independent pathway, and the microbial product, curdlan, via a PRR-dependent pathway, display convergence of signaling pathways on CREB in cDC2s, but not cDC1s. Consequently, IRF4 and KLF4 levels are repressed, resulting in a pro-Th17 phenotype of cDC2s (i.e., cDC17s). These results define a novel molecular event by which cDC2s undergo re-programming and hence affect Th cell differentiation, i.e., the induction of Th2 vs. Th17 bias (**Figures 1, 3 and Supp. Figure 3**).

Recent studies have mainly focused on the up-regulation of IRF4 expression. Here, we discovered that IRF4 expression is repressed by cAMP or curdlan signaling. DCs signaled in this manner skew the consequent T cell responses towards the Th17 effector lineage. In contrast, we previously found that when DCs signal through FcγRIII, IRF4 expression is up-regulated and this in turn leads those DCs to skew T cell responses towards a Th2 effector lineage (Williams et al., 2013). The current results indicate that cAMP-PKA-CREB signaling is a non-PRR pathway that reprograms DCs. Reduced cAMP signaling in DCs provokes Th2 differentiation (Lee et al., 2015) and increased cAMP signaling provokes Th17 differentiation (Datta et al., 2010). Furthermore, increasing the cAMP concentration in pro-Th2 cDC2s can switch them toward cDC17 phenotype. DCs thus appear to integrate stimuli from their extracellular microenvironment to control their IRF4 expression levels, which in turn, regulates the ensuing T cell response.

The effects of cAMP (i.e., CPT) were observed in two different types of DCs; splenic cDC2s and BM-APCs but not in cDC1s cells (**Figure 1 and Supp. Figure 1, 3**). The cAMP-regulated responses are achieved via the inhibition of *Irf4* and *Klf4,* TFs that are prerequisites for the pro-Th2 DC phenotype of cDC2(Gao et al., 2013; Tussiwand et al., 2015; Williams et al., 2013). cDC2s from *Gnas*^ΔCD11c^ mice (cells with prominently decreased cAMP signaling) have higher levels of *Irf4* and *Klf4* than do cDC2s from WT mice (**Supp. Figure 2**). The *Irf4* and *Klf4* transcript and IRF4 protein levels were inhibited by multiple cAMP agonists and by the PRR Dectin-1 activated by curdlan (cAMP-independent) in WT cDC2s, which suggests convergent signaling to inhibit *Irf4* and *Klf4* for the cDC17 phenotype (**Figure 3**). Based on previous data (Bedoui & Heath, 2015; Persson et al., 2013; Schlitzer et al., 2013), we were surprised that BM-APCs and cDC2s from two mouse strains in which IRF4 had been genetically-deleted provoked a strong Th17 response in the presence of very low IL-1 and IL-6 secretion, suggesting that this pathway of Th17 induction is independent of these cytokines (**Figure 4F-H**). Furthermore, curdlan provoked neither cAMP synthesis nor PKA phosphorylation, but activates CREB (Elcombe et al., 2013), possibly via MAPK signaling (Kim et al., 2016). *In vivo,* pro-Th17 DCs induction by cAMP-independent or PRR-dependent signaling pathways were confirmed by adoptive transfer of CPT- or curdlan-treated cDC2s to WT recipients, which resulted in an increased number of neutrophils and IL-17A production in the airway (**Figure 6, Supp. Figure 10**).

Collectively, our data indicate that: 1) inhibition of the pro-Th2 DC cDC2 phenotype by cAMP agonists promotes induction of the pro-Th17 DC cDC17 phenotype, and 2) reduction of IRF4 levels is necessary but not sufficient for induction of the pro-Th17 transcriptional program in DCs by both the PRR-dependent and cAMP-dependent signaling pathways. The results further indicate that the pro-Th17 DC phenotype induced by cAMP also requires IRF5. We therefore propose that the currently defined cDC2 subset can be redefined as two separate subpopulations: cDC17 (CD11c^+^, CD11b^+^, IRF4^-^ or IRF4^low^, IRF5^+^), and cDC2 (CD11c^+^, CD11b^+^, IRF4^+^, KLF4^+^).

Our findings indicate that endogenous IRF4 is a transcriptional repressor of a subset of cAMP-dependent genes in cDC2s, based on the following: 1) its loss causes up-regulation of cAMP-dependent gene expression, 2) the presence of an IRF4 binding site is highly predictive of a gene that is regulated by cAMP, and 3) an IRF4 binding site predicted that a gene increases (but not decreases) in *Irf4*^-/-^ cDC2s and decreases in *Irf4* restored cells. Overexpression of IRF4 restored the repression of these genes but also induced expression of genes not normally engaged by IRF4. The presence of a KLF4 binding site in the context of IRF4 over-expression was predictive of down-regulation of gene expression, consistent with the known role of KLF4 as a transcriptional repressor. IRF4 binding sites tended to be associated with enhancers as well as promoters and showed the three known IRF4 binding motifs (50% EICE, 23% AICE, 24% ISRE). While all three sites were associated with transcriptional activity, only the ISRE was associated with transcriptional repression. The AICE was most strongly associated with up-regulated genes by CPT, perhaps indicating a role for the bZIP binding partner BATF/JUN in this response. We therefore conclude that IRF4 is a major repressor of cAMP-stimulated gene expression in cDCs.

A consequence of this inverse relationship is that a large portion of cAMP-stimulated genes is, in fact, indirectly regulated through suppression of *Irf4* expression in these cells. The mechanism involved in the repression of *Irf4* is unknown. Intriguingly, the orphan nuclear receptor NR4A1 (NUR77) represses *Irf4* expression in CD8^+^ T cells by binding to multiple sites in the *Irf4* promoter (Nowyhed, Huynh, Thomas, Blatchley, & Hedrick, 2015). In addition, OVA-challenged *Nr4a1* KO mice have increased airway inflammation, and elevated Th2 cytokines and eosinophil count in BAL fluid (Kurakula et al., 2015). In Leydig cells, cAMP activates expression of *Nr4a* (Mori Sequeiros Garcia, Gorostizaga, Brion, Gonzalez-Calvar, & Paz, 2015) but we find that CPT treatment represses *Nr4a1* expression in cDC2s so this is unlikely to be the mechanism. In contrast, the related family member NR4A3 (NOR1) stimulates *Irf4* expression in BMDC (Nagaoka et al., 2017), so NR4A family members may regulate *Irf4* expression through differential recruitment of co-repressor and co-activators. Among other transcriptional regulators, NF-κB, NFAT, SP1 and STAT4 activate *Irf4* expression (Bat-Erdene et al., 2016; Boddicker et al., 2015; Lehtonen et al., 2005; Sharma et al., 2002). How cAMP signaling interacts with these pathways in cDC2s is not known but VIP/cAMP signaling interferes with IL-12/JAK2/STAT4 and NF-κB signaling in T cells and macrophages (Leceta et al., 2000; Liu, Yen, & Ganea, 2007). The regulation could also be epigenetic as expression of both *Nr4a1* and *Irf4* are repressed by HDAC7 in naïve T cells (Myers et al., 2017).

Reduction of IRF4 expression by cAMP elevation or genetic knockout induced a number of genes encoding secreted proteins in cDC2s, some of which have roles in Th differentiation and immune regulation. The *Ebi3* gene encodes IL-39 (an IL-12 family member), which has been implicated in autoimmunity (X. Wang et al., 2018), repression of immune responses to *T. cruzi* (Bohme et al., 2016), and Th17-mediated immunity to *L. monocytogenes* (Chung et al., 2013). *Il1rn-*deficient mice develop autoimmune IL-17A-mediated arthritis (Ikeda et al., 2014; Rogier et al., 2017), *Mmp14* has been implicated in DC podosome protrusion and invasion and antigen presentation (Baranov et al., 2014; Gawden-Bone et al., 2010), and platelet factor 4 inhibits CD1a expression on DC and limits Th17 differentiation (Gerdes et al., 2011; Xia & Kao, 2003). Tissue plasminogen activator has not been directly implicated but an inhibitor of plasminogen activator inhibitor-1 reduces Th2 cytokines and bronchial eosinophils in HDM-induced asthma in mice (Tezuka et al., 2015). Lung epithelial cells from *Thbs4*-/-mice exhibit decreased adhesion, migration and proliferation (Muppala et al., 2015) and variants in *THBS4* are associated with altered lung function (Panasevich et al., 2013). Many of these secreted, cAMP-and IRF4-dependent genes, such as Lipocalin-2 (LCN2) (Hau et al., 2016) and Leucine-rich alpha 2 glycoprotein (LRG) (Urushima et al., 2017) may have functional roles in Th17 differentiation (**Figure 5C**), perhaps contributing to the independence of Th17 induction from IL-1 and IL-6 (**Figure 4F-H**) by this pathway.

Microbial stimulation most likely does not underlie the cAMP-mediated Th2 and Th17 responses for several reasons: 1) the *in vivo* Th2 and Th17 differentiation were generated *in vitro* in the presence of antibiotics, 2) the KO mice were co-housed with *fl/fl* littermates, which showed no Th bias, 3) *fl/fl* BM-APCs and cDC2s display a WT phenotype under the same conditions and 4) WT and *Gnas*Δ^CD11c^ BM-APCs and cDC2 could be reprogrammed *in vitro* to a pro-Th17 phenotype by multiple cAMP-elevating agents (in the presence of antibiotics). Together, these factors strongly suggest the lack of a major impact of the microbiota or their products on the Th bias observed in our experiments.

As part of homeostatic regulation, cAMP is regulated by GPCRs that recognize host-derived ligands, such as neurotransmitters, hormones, chemokines, metabolites (including potentially microbial-derived metabolites, such as fatty acids, lipid mediators, nucleosides/nucleotides (Marinissen & Gutkind, 2001)), complement cleavage fragments, cholera (Datta et al., 2010), pertussis and heat-labile toxins (Bharati & Ganguly, 2011), and bacterial formylated peptides (Cattaneo, Parisi, & Ammendola, 2013). Thus, GPCRs that alter cellular cAMP concentrations likely have a role in innate immunity and consequently, in Th differentiation. However, the current studies do not reveal the physiologically most important GPCR(s) that mediate this process.

GPCR pathway drugs (Sriram & Insel, 2018) and others that regulate cAMP levels, including cyclic nucleotide phosphodiesterase (PDE) inhibitors, have the potential to alter Th2/Th17 bias and potentially may cause immunological side effects, or alter diseases that involve Th2 or Th17 responses. Drugs that increase cAMP levels in DCs perhaps may facilitate recovery from bacterial infections such as *K. pneumonia* and *P. aeruginosa,* and fungal infections, e.g., *candida albicans* (Kumar, Chen, & Kolls, 2013), for which Th17 response is beneficial. On the other hand, drugs that decrease cAMP levels in DCs and therefore enhance Th2 immunity, might facilitate treatment of helminth infections. The inhibition of IRF4 by agonists that increase intracellular cAMP may also have implications for other disease settings. IRF4 is a key regulator at multiple steps in B-cell differentiation and development (Klein et al., 2006; Ochiai et al., 2013; Sciammas et al., 2006) and is an oncogene in multiple myeloma (Shaffer et al., 2009), to a lesser degree in Hodgkin and non-Hodgkin lymphomas as well as in chronic lymphocytic leukemia (Boddicker et al., 2015; Shaffer et al., 2008; Shukla, Ma, Hardy, Joshi, & Lu, 2013). The current findings suggest that new (or perhaps repurposed) agonists of Gαs-linked GPCR drugs might be beneficial for the inhibition of IRF4 in B-cell derived malignancies.

In summary, the current studies revealed that both cAMP-PKA-CREB and PRR-CREB signaling reprogram DCs by regulating a set of TFs that control cDC2 and cDC17 phenotypes and subsequent Th2 or Th17 bias. These findings thus demonstrate a previously unappreciated PRR-independent, transcriptional mechanism for Th bias by DCs and suggest new therapeutic approaches to control immune responses.

## Materials and Methods

### Mice and Cells

C57Bl/6 (B6), CD11c-Cre transgenic, OT2 (B6), IL17A-eGFP and loxP-flanked *Irf4* (*Irf4*^fl/fl^) mice were purchased from The Jackson Laboratory (Bar Harbor, ME). *Gnas*^ΔCD11c^ mice were produced in our laboratory by crossing of loxP-flanked *Gnas* mice (a kind gift from Lee Weinstein, NIH) with CD11c-Cre mice as described previously (Lee et al., 2015). To generate *Irf4*-deficient CD11c cells, *Irf4*^fl/fl^ mice were crossed to CD11c-Cre mice for at least 4 generations. Cre-mediated GFP expression was used to trace *Irf4* ablation with other markers (CD11c, CD11b and CD8α) in splenocytes (Klein et al., 2006; Mildner & Jung, 2014). More than 96% of isolated cDC2s were GFP positive. The fl/fl littermates (Cre^-^ *Irf4*^fl/fl^) were used as controls. IL4-eGFP reporter (4Get) mice were originally made by Dr. R. Locksley (University of California San Francisco) (Mohrs, Shinkai, Mohrs, & Locksley, 2001) and were a gift from Dr. M. Kronenberg (LAI, San Diego, CA). IL17A-eGFP mice were bred to OT2 mice to yield 4Get/OT2 and IL17A-eGFP/OT2 mice, respectively. All mice were kept in a specific pathogen-free (SPF) facility. *Irf4*-inducible (*Irf4^-/-^*) mice (Ochiai et al., 2013) were bred by Dr. R. Sciammas (University of California, Davis). *Irf5*^-/-^ mice were provided by Dr. I. R. Rifkin (Boston University) (Takaoka et al., 2005; Watkins et al., 2015). IL17A^-/-^ mice were obtained from Dr. Y. Iwakura (The University of Tokyo, Japan) (Nakae et al., 2002).

OT2 T cells were isolated by magnetic beads (EasySep™ Mouse Naïve CD4^+^ T Cell Isolation Kit, StemCell Technologies) from a single cell suspension of splenocytes. Bone marrow (BM) cells were cultured in the presence of GM-CSF (10 ng/ml) for 7 days. Floating cells from the BM culture were applied to FACS-sorting and CD11c^+^CD135^+^ BM cells (BM-APCs) were used for co-culture with naïve OT2 T cells and transcription factor (TF) analysis by qPCR. cDC2 cells were isolated from single cell suspension of spleens. CD11c^+^CD11b^+^CD8α^-^ splenocytes (Mildner & Jung, 2014; Worbs, Hammerschmidt, & Forster, 2017) were isolated by FACS sorting and applied to co-culture and TFs analysis. cDC1 were isolated by FACS sorting of CD11c^+^CD11b^-^CD8α^+^ splenocytes and subjected to the same analysis described for cDC2.

### Reagents

Reagents obtained are as follows: 8-(4-Chlorophenylthio) adenosine 3′,5′-cyclic monophosphate sodium salt (8-CPT-cAMP), forskolin, prostaglandin E2 (PGE2), pertussis toxin (PTX), dexamethasone, Phorbol 12-myristate 13-acetate (PMA), ionomycin, alum, doxycycline and CREB inhibitor (666-15) were from Sigma-Aldrich; Cholera toxin was from list biological laboratories; Ovalbumin (purified) was from Worthington Biochemical; MHC class II OVA peptide (OVA323-339: ISQAVHAAHAEINEAGR) was from GenScript; HDM was from Greer; anti-mouse CD3e (clone 2C11) antibody and anti-CD28 antibody were from BioXcell; Rp-cAMP was from Biolog; rolipram (PDE4 inhibitor) and EPAC inhibitor (CE3F4) were from Tocris. Curdlan was from Wako Chemicals; All the antibodies used for FACS were purchased from BD Pharmingen, eBiosciences or BioLegend.

### OVA-specific immune responses in the cDC-OT2 co-culture

OVA-specific CD4^+^ T cell response was performed using the DC-OT2 co-culture system as described (Lee et al., 2015). Briefly, CD11c^+^CD11b^+^CD8α^-^ splenocyte (cDC2s), CD11c^+^CD11b^-^ CD8α^+^ splenocyte (cDC1s) or CD11c^+^CD135^+^ BM cells (BM-APCs) were isolated by FACS sorting as described above. cDC2s and cDC1s were loaded with OVA peptide (1 μg/ml) 2 hr before T cell addition and BM-APCs were cultured in complete RPMI 1640 containing OVA protein (100 μg/ml) for 16hr. Various cAMP agonists indicated in the figures were added in the culture 16 hr before T cell engagement. Specially, curdlan (10 μg/ml) was treated for 24 hr before co-culture. For the IRF4 restoration in IRF4^-/-ind^ cDC2s, 200 ng/mL doxycycline was added for 12 hr before co-culture. OT2 T cells were co-cultured with cDC2 (3×10^5^ cells) at 1:2 ratio in a round-bottom 96-well plate or with BM-APC (5×10^5^ cells) at 1:1 ratio in a 24-well plate in the serum-free culture medium supplemented with albumin. After 3 days of co-culture, OT2 T cells were stimulated with plate-bound anti-CD3/28 antibodies for 24hr and then used by ELISA to measure cytokines levels or stimulated with PMA and ionomycin for 3hr for the T cell lineage marker (qPCR).

### Switch from memory Th2 to Th17 (Fate mapping)

Memory Th2 cells (TEM, CD44^+^CD62L^low^ or TCM, CD44^+^CD62L^high^) were generated by co-culturing of naïve IL-17GFP CD4^+^ OT2 cells with BM-APC from *Gnas*^ΔCD11c^ mice for 3 days and then T1/ST2^+^ cells were sorted by FACS. T1/ST2^+^ cells were used for 2^nd^ co-culture with OVA peptide (1 μg/ml) loaded-CPT (50 μM) or Cholera toxin (1 μg/ml) treated WT cDC2s. After 3 days of 2^nd^ co-culture, IL-17GFP CD4^+^ OT2 T cells were stimulated with plate-bound anti-CD3/28 antibodies for 24hr and then used for ELISA to measure cytokines levels or stimulated with PMA (50 ng/ml) and ionomycin (1 µM) for 6hr for the GFP expression measurement by FACS.

### Human DC-like cells differentiation

Human myeloid leukemia cell line MUTZ-3 were acquired from Dr. Martin L. Yarmush (Rutgers University School of Engineering, Piscataway, New Jersey). MUTZ-3 cells were maintained and differentiated into DC-like cells as described previously (Koria, Bhushan, Irimia, & Yarmush, 2012). Briefly, MUTZ-3 cells were cultured in α-MEM (Invitrogen, Carlsbad, CA, USA) supplemented with 20% Fetal bovine serum, 50 µM β-mercaptoethanol and 10% 5637 cell-conditioned media. For generation of DC-like cells, MUTZ cells (10^5^ cells/ml, 2 ml medium/well) were cultured in growth media supplemented with GM-CSF (100 ng/ml, BioLegend, San Diego), TGF-β (10 ng/ml, R&D systems) and TNF-α (2.5 ng/ml, R&D systems) for 7 days. The human monocytic leukemia cell line, THP-1 was from ATCC. As described previously (Berges et al., 2005), THP-1 cell line was maintained in RPMI 1640 supplemented with 10% FCS at a concentration of 2 × 10^5^ cells/ml. To induce differentiation, rhIL-4 (100 ng/ml) and rhGM-CSF (100 ng/ml) were added. Cells were cultured for 5 days to acquire the DC-like cell phenotype. Medium was exchanged every 2 days with fresh cytokine-supplemented medium. The human leukemia cell line, HL-60 was from ATCC. HL-60 cells were cultured in RPMI 1640 medium containing 10% FCS. For DC-like differentiation, HL-60 cells were incubated in the culture media together with calcium ionophore A23187 (180 ng/ml) and rhGM-CSF (100 ng/ml) for 24 hr, as described (Yang, Li, & Zhao, 2007).

### ELISA measurement of cytokines

Cytokine levels in the supernatant were determined using ELISA kits for IL-4, IL-5, IL-10, IFN-γ and IL-17A (eBioscience, La Jolla, CA) following the manufacturer’s instructions.

### Flow cytometry and intracellular staining

The data were acquired by a C6 Accuri flow cytometer (BD Biosciences) and analyzed by FlowJo Software. For the staining of surface molecule, cells were washed with FACS buffer (2% FCS containing PBS) and incubated with the indicated antibodies on ice for 30 min. After two times of wash with FACS buffer, cells were used for analysis. For IRF4, IRF5 and IRF8 intracellular staining, surface marker stained cells were fixed and permeabilized using Cytofix/Cytoperm™ (BD Biosciences) and stained with Ab for 30 min. After 3 times of wash with permeabilization buffer, the intracellular levels of IRFs were analyzed in the different DC subsets. For the detection of eGFP^+^ cells, harvested CD4^+^ T cells or lung single cells were stimulated with PMA (50 ng/ml) and ionomycin (1 µM) in the presence of GolgiStop (BD Pharmingen) for 6hr before FACS analysis.

### Adoptive transfer of cDC2s

Adoptive transfer model was initially described by Lambrecht B. et al (Lambrecht, Pauwels, & Fazekas De St Groth, 2000). Briefly, splenic cDC2s from WT donors were pulsed with HDM (50 μg/ml) or OVA (100 μg/ml), with and without CPT or Curdlan. After 24h incubation, the cDC2s were washed with PBS three times, re-suspended in PBS, and transferred into anesthetized WT or IL-17eGFP recipient mice i.n (5 × 10^5^ cells in 20 µl PBS) on day 0 and 14 (Lee et al., 2015; Machida et al., 2004). Mice were challenged by 12.5 µg HDM, or 25 µg OVA i.n. on day 19, 20 and 21. One day after the last challenge, mice were analyzed for airway hyper-responsiveness (AHR) to methacholine (MCh). Bronchoalveolar lavage (BAL) fluid was collected for cellular composition (light microscopy) and cytokine analysis. For cytokine analysis from lung tissue, lung single cell suspension was prepared from three lobes from each mouse as described (Doherty et al., 2011; Lee et al., 2015) and stimulated with HDM (50 μg/ml) or OVA (200 μg/ml) for 3 days. Supernatant from the culture was applied to cytokine analysis by ELISA.

### Quantitative PCR analysis

Isolation of RNA was carried out using an RNA purification Kit (Thermo Fisher Scientific) according to the manufacturer’s instructions. The cDNA was synthesized using Superscript III First-Strand system (Invitrogen). Quantitative PCR analysis was performed as described previously (Lee et al., 2015). SYBR Green PCR Master Mix was used for real-time PCR (Thermo Fisher Scientific). Samples were run in triplicate and normalized by GAPDH. PCR for *Irf5* was performed using Taqman primers according to the manufacturer’s instructions. Primer sequences are listed in previous study (Lee et al., 2015) and additional sequences are *Irf4*: F-AGATTCCAGGTGACTCTGTG, R-CTGCCCTGTCAGAGTATTTC, *Irf8*: F-CGCTGTAGGAAAAGCAGACC, R-CCTCCAACAACACAGGGAGT, *Klf4*: F-CTGAACAGCAGGGACTGTCA, R-GTGTGGGTGGCTGTTCTTTT, *Crem*: F-GCTGAGGCTGATGAAAAACA, R-GCCACACGATTTTCAAGACA, Il-23p19: F-TCCGTTCCAAGATCCTTCG, R-GAACCTGGGCATCCTTAAGC.

### Expression profiling and genomic analysis

Total RNA was isolated from splenic cDC2 cells using RNeasy columns (Qiagen, Germantown, MD). Non-stranded sequencing libraries were prepared using Illumina TruSeq RNA library kits and sequenced on an Illumina Hi Seq2500 using a SE70 protocol (Illumina, San Diego, CA). Raw sequenced reads were trimmed for adapter and bar-codes sequences and quality assessed using FastQC (http://www.bioinformatics.babraham.ac.uk/projects/fastqc/). Reads were aligned to the mm10 genome using STAR (Dobin et al., 2013) with mgcGene annotations (https://genome.ucsc.edu/cgi-bin/hgTables). Differential expression was determined using DESeq2 (Anders & Huber, 2010) in SeqMonk (http://www.bioinformatics.babraham.ac.uk/projects/seqmonk/). ChIPseq and ATACseq reads were identified using the Cistrome Data Browser (http://cistrome.org/db/#/) then raw reads were downloaded from the Sequence Read Archive (https://www.ncbi.nlm.nih.gov/sra) using sratools (https://ncbi.github.io/sra-tools/). The reads were aligned with STAR then binding peaks identified using the HOMER suite (http://homer.ucsd.edu/). The binding-expression predictions were run with BETA (S. Wang et al., 2013). Histone modification co-localization, the transcription factor binding metagene analysis and the de novo motif identification were performed using HOMER. RNAseq, ChIPseq, ATACseq and histone modifications were visualized in IGV (Robinson et al., 2011). Heatmaps were generated using heatmap.2 in R. Transcriptional network and pathway enrichment analysis were performed in MetaCore (Genego, Clarivate Analytics, Philadelphia, PA).

### Statistical analysis

Student’s t-tests were used to analyze data sets with two groups (GraphPad Prism software). One- or two-way ANOVA were used for multiple groups. All data are represented as mean ± s.e.m unless indicated otherwise. *p*-values<0.05 were considered significant.

### Study Approval

All the experimental procedures were approved by the UCSD-IACUC.

## Author contributions

Conception: J.L., N.J.G.W. and E.R.; discussion: J.L., J.M.G.-N., P.A.I, M.C., A.T., R.C., N.J.G.W. and E.R.; research design and experimentation: J.L, J.Z, Y.-J.C., J.H.K, C.M.K., J.M.G.-N. and D.S.H.; data analysis: J.L, J.Z, Y.-J.C., J.H.K, C.M.K., J.M.G.-N., P.A.I, M.C., K.Y., I.R.R., D.B. R.C., N.J.G.W. and E.R.; resource assistance and clinical samples: D. S. H., B. N., M.C., K.Y., I.R.R., D.B. R.C. and N.J.G.W.; writing of draft and editing: J.L., P.A.I., N.J.G.W. and E.R.

## Acknowledgements

We would like to acknowledge assistance from the Genomic and the Histology Shared Resources that are supported by the Moores Cancer Center CCSG Grant (NIH CA023100), UCSD/UCLA Diabetes Research Center Grant (NIH DK063491), Dr. Nissi Varki for her invaluable help in the assessment of mouse pathology and James Lee for assistance with cell preparation. This paper is dedicated to the late Dr. Tim Bigby, a pulmonologist who provided invaluable insights and advice.

## Competing interests

The authors declare no competing interests.

## Data and materials availability

All data supporting the findings of this study are available within the paper or in the supplementary materials.

---

**Supplemental Figure 1.**
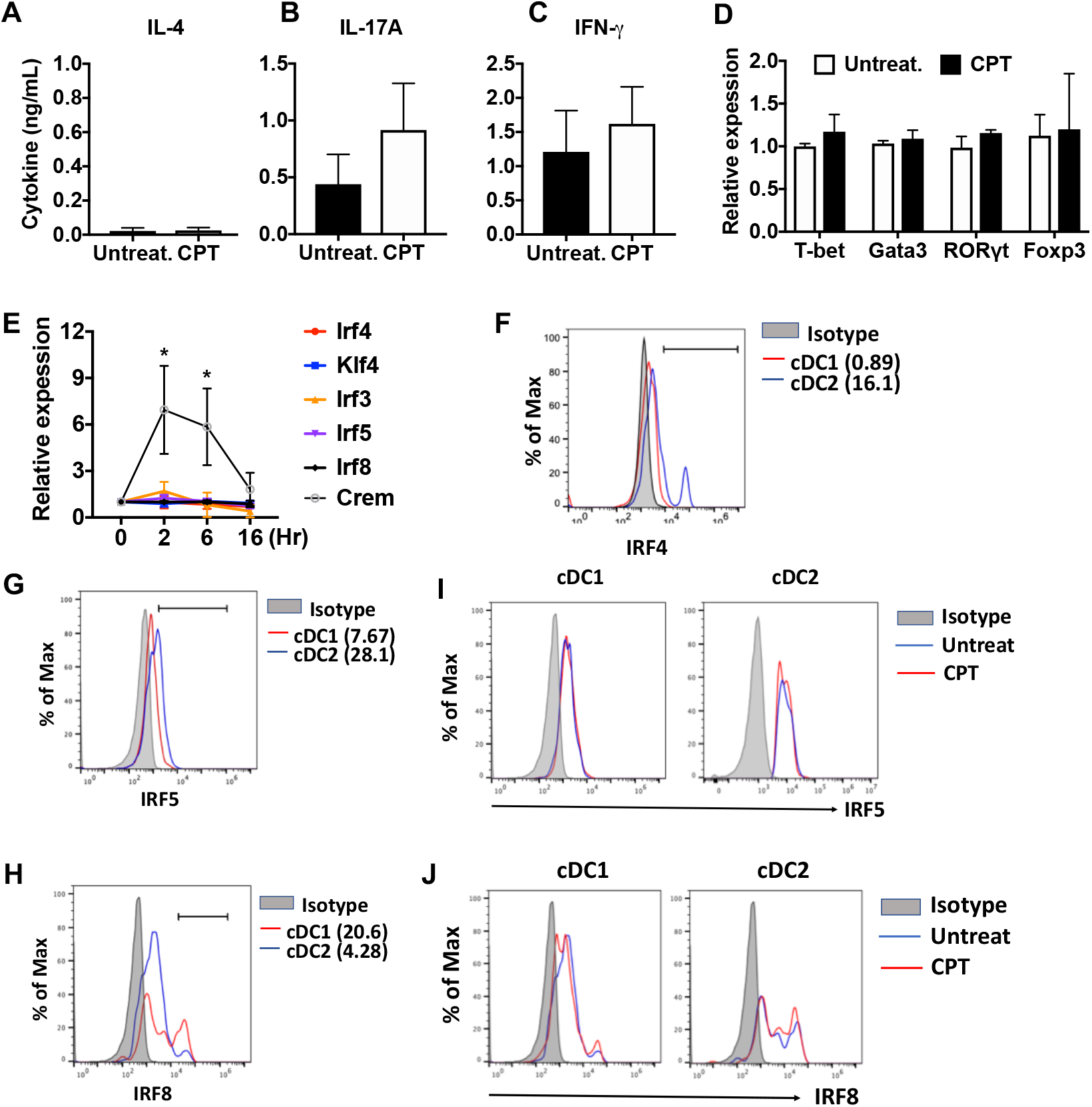
cAMP signaling in cDC1s does not affect the expression of IRF and subsequent T cell differentiation. **(A-C)** IL-4, IL-17A and IFN-γ levels from anti-CD3/28 Ab-stimulated OT2 cells co-cultured with cDC1s (CD11c^+^CD11b^-^CD8α^+^) cells from WT mice pretreated with or without CPT. **(D)** QPCR analysis of lineage commitment factors in OT2 cells co-cultured with WT cDC1s in the presence of CPT. **(E)** Relative expression of TFs in WT cDC1s treated with CPT for indicated time. **(F)** IRF4, **(G)** IRF5 and **(H)** IRF8 expression in WT cDC1s and cDC2s. **(I)** IRF5 and **(J)** IRF8 expression in cDC1s and cDC2s after CPT treatment for 48 hr. Data are mean ± s.e.m, *n*=3 in each group; * *p*<0.05.

**Supplemental Figure 2.**
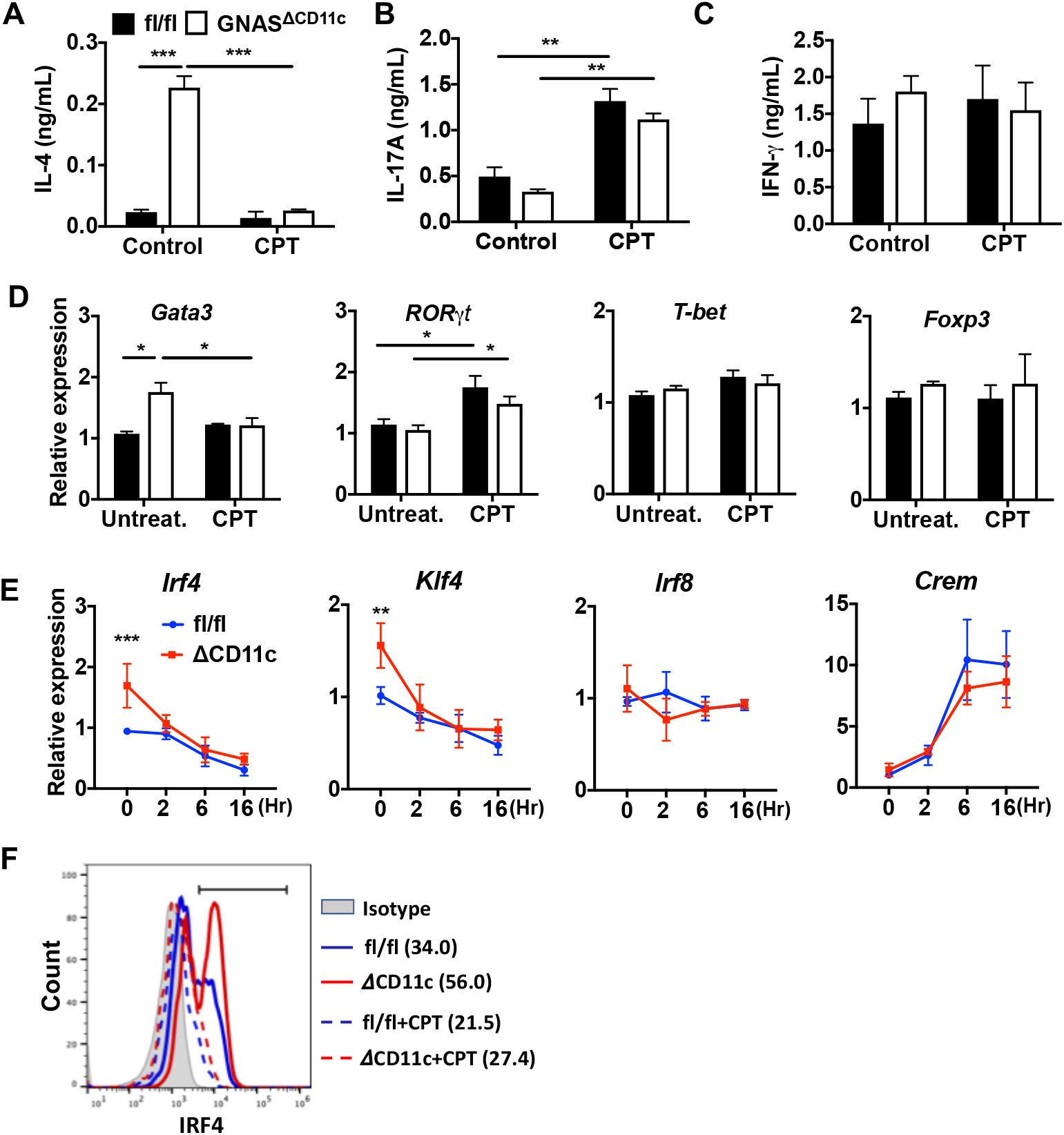
cAMP signaling switches a pro-Th2 *Gnas*^ΔCD11c^ to a pro-Th17 phenotype. **(A-C)** IL-4, IL-17A and IFN-γ levels from anti-CD3/28 Ab-stimulated OT2 cells co-cultured with splenic cDC2s from fl/fl and *Gnas*^ΔCD11c^ mice treated in the absence or presence of CPT. **(D)** QPCR analysis of lineage commitment factors in the isolated OT2 cells co-cultured with fl/fl or *Gnas*^ΔCD11c^ cDC2s. **(E)** QPCR of TFs in CPT-treated fl/fl or *Gnas*^ΔCD11c^ cDC2s. Two-way ANOVA with Sidak’s multiple comparisons test; ** p<0.01, ***p<0.001. Effect of CPT treatment; *Irf4* (*p*<0.001), *Klf4* (*p*<0.001) and *Crem* (*p*<0.001). Effect of strain: *Irf4* (*p*<0.001). **(F)** Intracellular staining (FACS) of IRF4 in untreated and CPT-treated *Gnas*^fl/fl^ and *Gnas*^ΔCD11c^ cDC2s for 48 hr. Data are mean ± s.e.m, *n*=3 in each group; * *p*<0.05, ** *p*<0.01, \*\*\**p*<0.001.

**Supplemental Figure 3.**
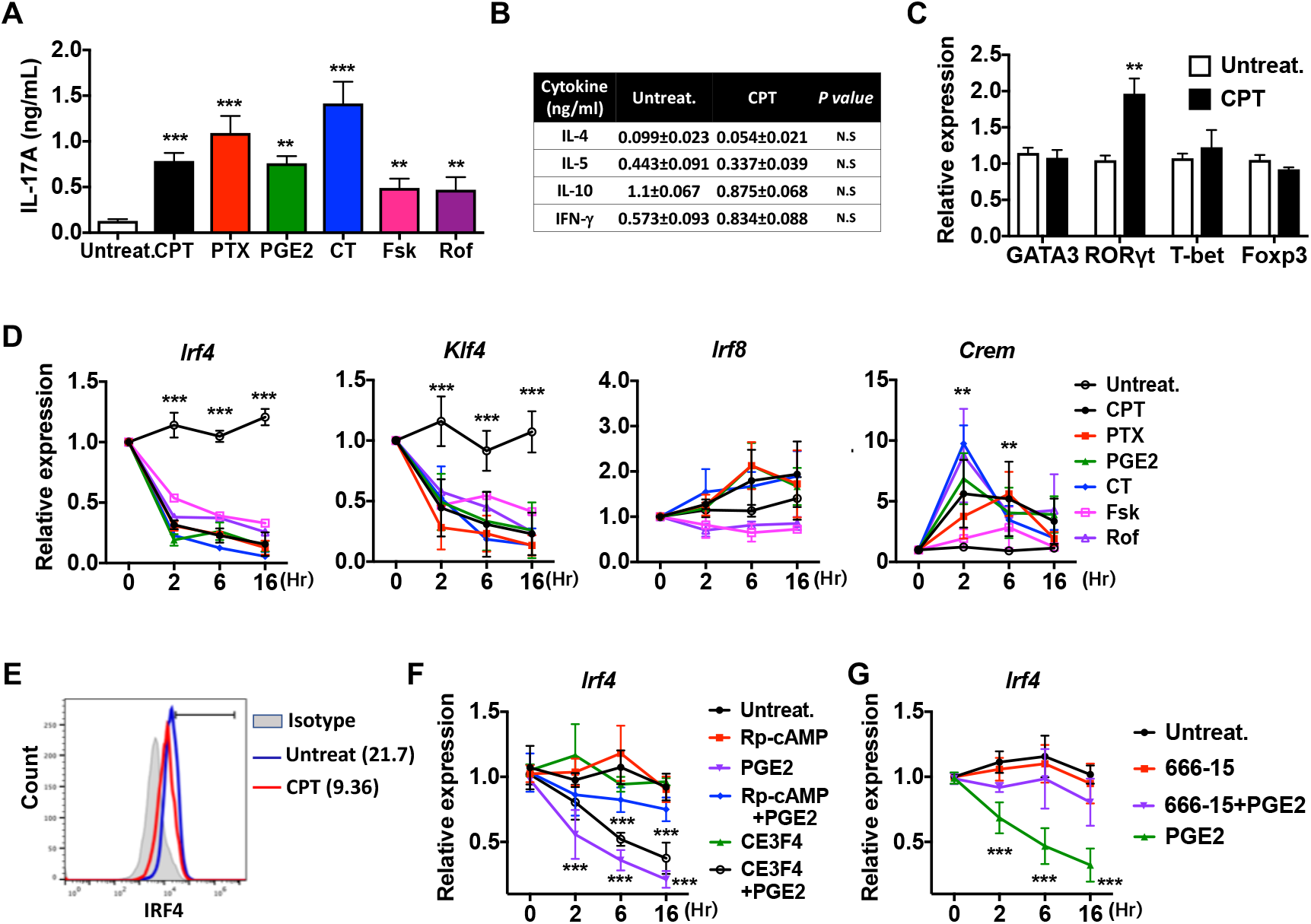
Induction of pro-Th17 BM-APCs and altered transcriptional program by cAMP agonists. **(A)** IL-17A levels from anti-CD3/28 Ab-stimulated OT2 cells co-cultured with BM-APCs from WT mice treated with various cAMP agonists: CPT (50 μM), pertussis toxin (PTX, which inhibits Gαi, 100 ng/ml), prostaglandin E_2_ (PGE_2_, 1µM), cholera toxin (CT, an activator of Gαs, 1 µg/ml), forskolin (Fsk, an activator of AC, 10 µM), rolipram (Rol, PDE4 inhibitor, 10 µM). **(B)** Levels of the indicated cytokines and **(C)** mRNA expression of lineage commitment factors in OT2 cells co-cultured with WT BM-APCs treated with CPT. **(D)** QPCR of TFs in the WT BM-APCs treated with the indicated cAMP agonists. Two-way ANOVA with Sidak’s multiple comparisons test; *n*=3 in each group, different from untreated in the CPT-treated group; * *p*<0.05, ** *p*<0.01, \*\*\**p*<0.001. Effect of treatment; *Irf4* (*p*<0.001), *Klf4* (*p*<0.001), *Irf8* (*p*<0.001) and *Crem* (*p*< 0.001). **(E)** Intracellular staining of IRF4 in CPT-treated WT BM-APCs for 48 h. **(F, G)** QPCR of *Irf4* in the WT BM-APCs treated with PGE_2_ in the presence of inhibitors of PKA (Rp-cAMP, 50 µM), EPAC (CE3F5, 50 µM), or CREB (666-15, 1 µM). Two-way ANOVA with Sidak’s multiple comparisons test; *n*=3 in each group, different from untreated; \*\*\**p*<0.001.

**Supplemental Figure 4.**
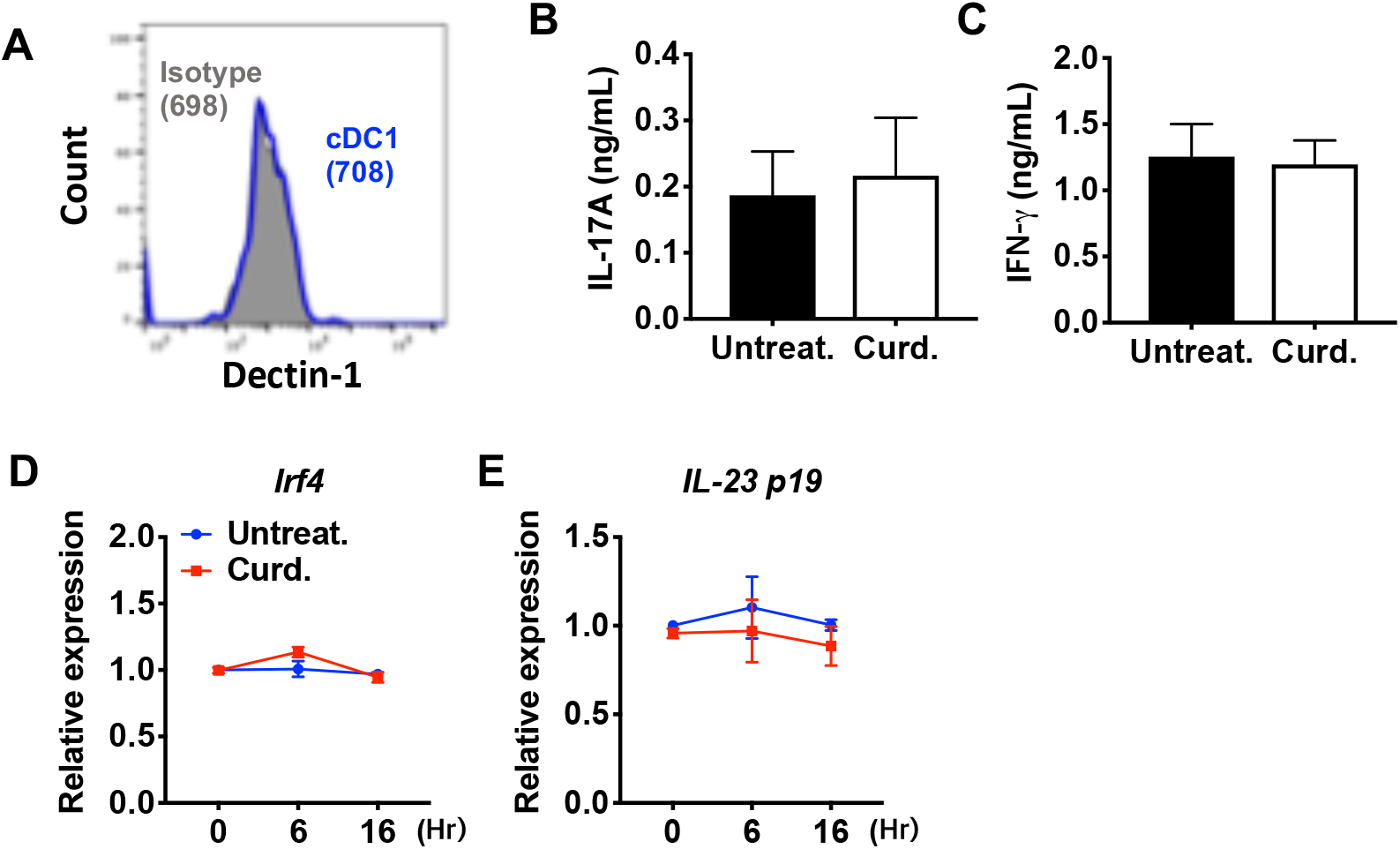
Curdlan does not affect the expression of *Irf4* and subsequent T cell differentiation in cDC1s. **(A)** Expression of Dectin-1 on WT cDC1s. Numbers indicate mean fluorescence intensity (GeoMFI). **(B)** IL-17A and **(C)** IFN-γ levels from anti-CD3/28 Ab-stimulated OT2 cells co-cultured with cDC1s from WT mice pretreated with or without curdlan. Relative expression of **(D)** *Irf4* and **(E)** IL-23 p19 mRNA in WT cDC1s treated with curdlan for the indicated time points. Data are mean ± s.e.m, *n*=3 in each group.

**Supplemental Figure 5.**
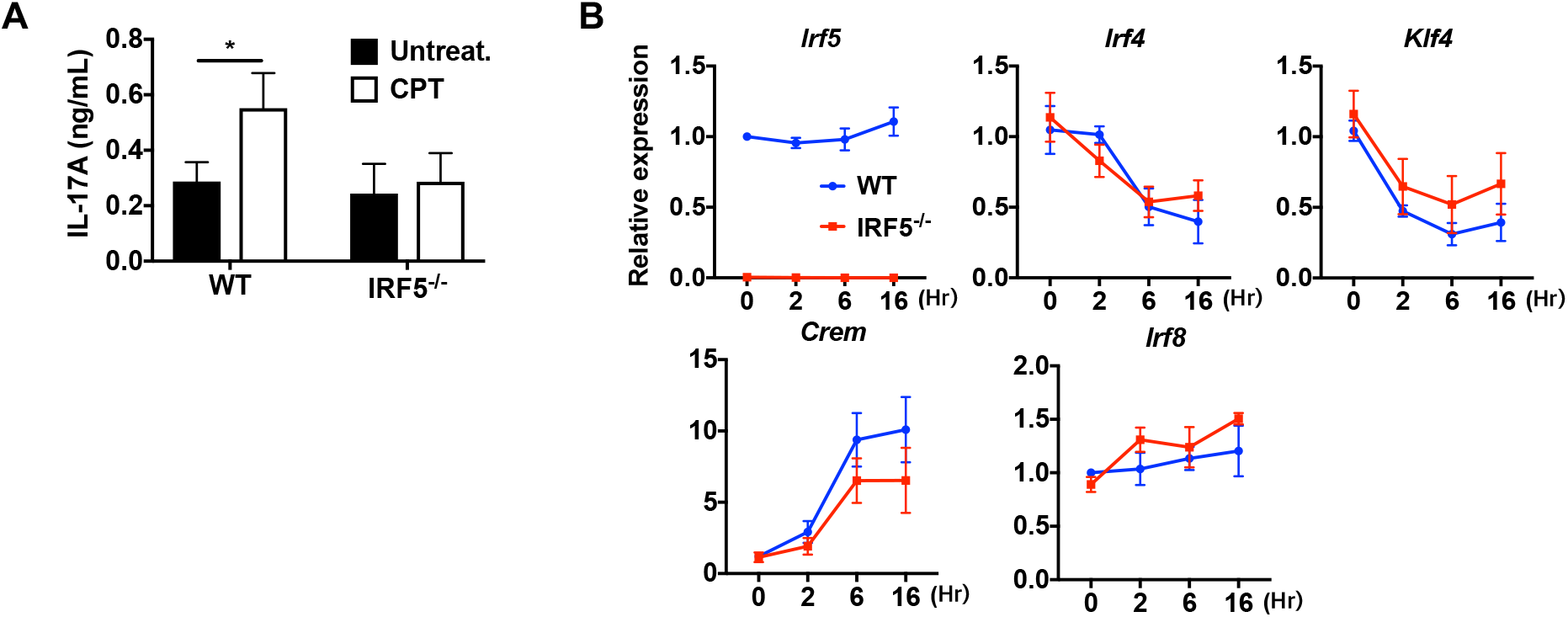
Loss of IRF5 in cDC2s inhibited CPT induced pro-Th17 phenotype. **(A)** IL-17A levels from anti-CD3/28 Ab-stimulated OT2 cells co-cultured with CPT-treated cDC2s from WT and *Irf5*^-/-^ mice. **(B)** QPCR of TFs in CPT-treated WT and *Irf5*^-/-^ cDC2s. Data are mean ± s.e.m, *n*=3 in each group; * *p*<0.05.

**Supplemental Figure 6.**
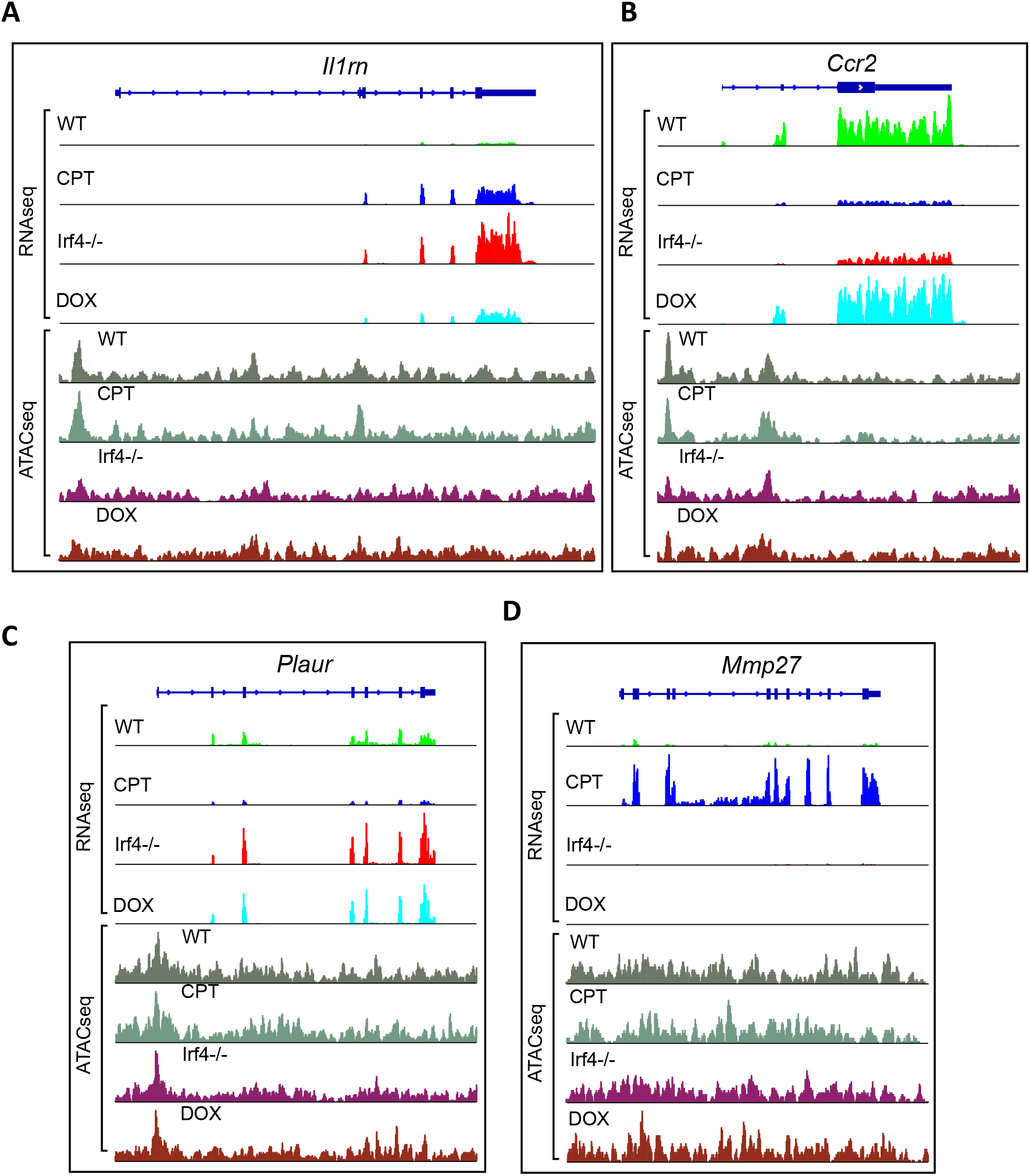
Examples of genes from transcriptional clusters. **(A-D)** Aligned RNAseq reads for the *Il1rn, Ccr2, Plaur* and *Mmp27* genes. Gene structure is indicated above in dark blue. Reads from untreated WT cells in green (WT), CPT-treated WT cells are in blue (CPT), *Irf4^-/-^* cDC2s in red (Irf4^-/-^) and Dox-induced *Irf4*^-/-^ cells in cyan (DOX). Chromatin accessibility was measured using ATACseq on the four groups. ATACseq peaks are shown below the RNAseq reads. Untreated cells are shown in gray (WT), CPT-treated cells in Gray-green, *Irf4*^-/-^ cells in purple (Irf4-/-) and Dox-induced *Irf4*^-/-^ cells in brown (DOX).

**Supplemental Figure 7.**
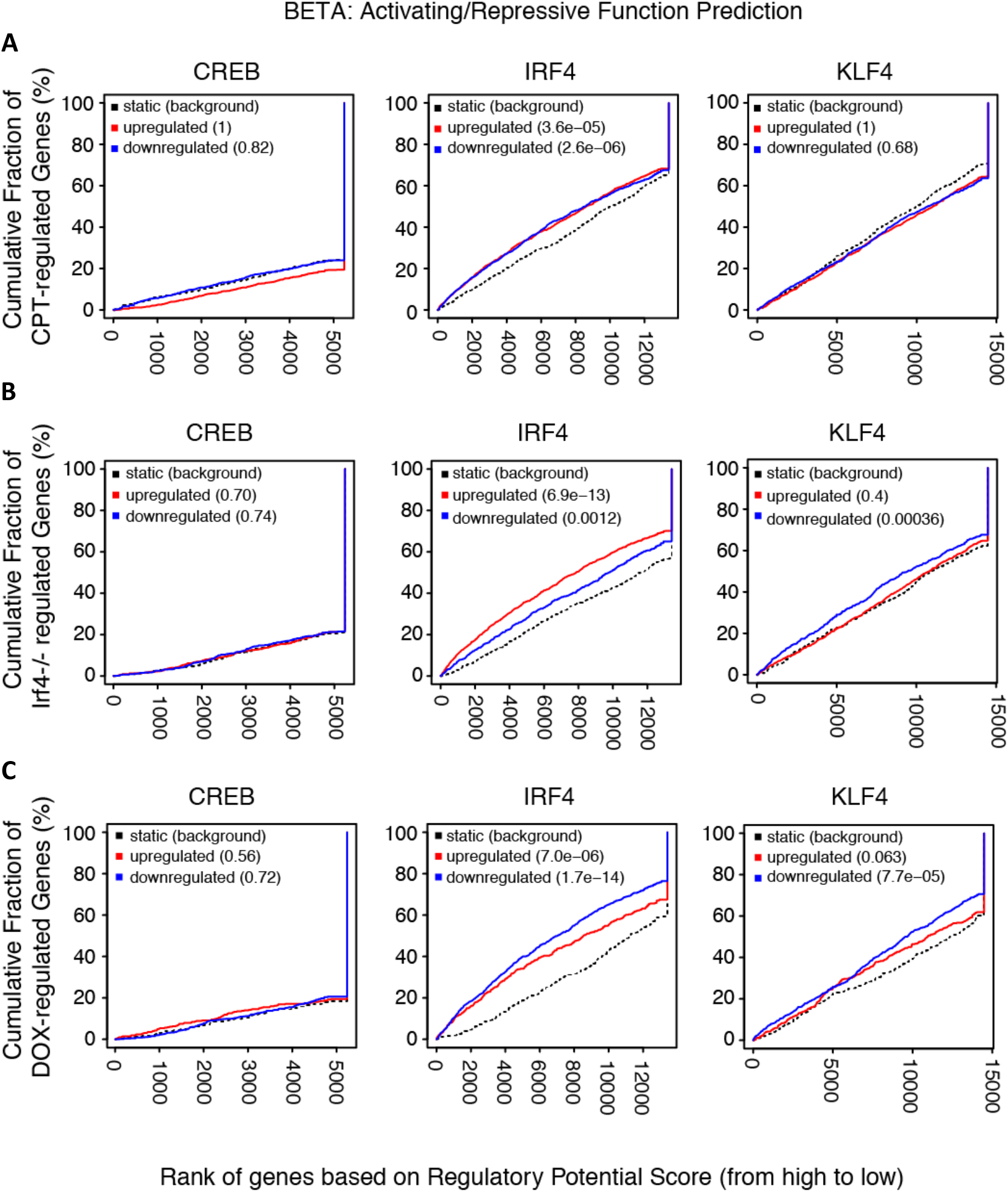
Transcriptional activation/repression prediction analysis. **(A-C)** The graphs show the cumulative fraction of regulated genes plotted against rank of genes on regulatory score potential using the BETA approach. Dotted black lines show expected background for no predictive value. Red curves indicate gene activation, blue curves indicate gene repression. p-values indicate significant difference between actual and expected curves for up-regulated and down-regulated genes in the CPT, *Irf4*^-/-^ and Dox datasets.

**Supplemental Figure 8.**
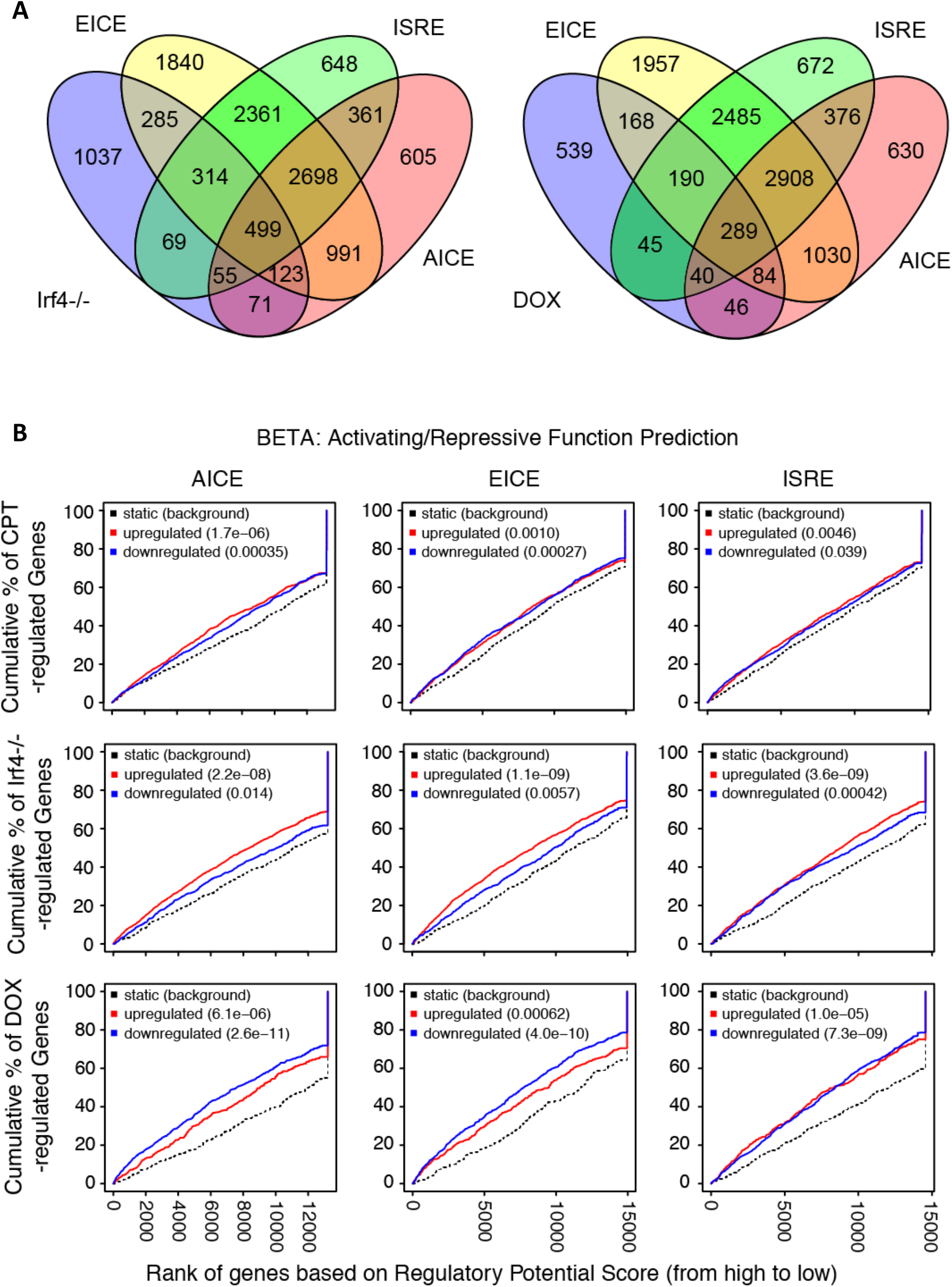
Prevalence and predictive value of IRF4 binding site subpopulations. **(A)** Venn diagrams showing overlap of EICE, AICE and ISRE motifs in genes from *Irf4^-/-^* cDC2s and from *Irf4-inducible* (*Irf4^-/-^*) cDC2s treated with Dox. **(B)** Prediction of CPT, *Irf4^-/-^* and Dox responsiveness with three IRF4 binding motifs. The graphs show the cumulative fraction of regulated genes plotted against rank of genes on regulatory score potential using the BETA approach. Dotted black lines show expected background for no predictive value. Red curves indicate gene activation, blue curves indicate gene repression. p-values indicate significant difference between actual and expected curves for up-regulated and down-regulated genes in the CPT, *Irf4*^-/-^ and Dox datasets.

**Supplemental Figure 9.**
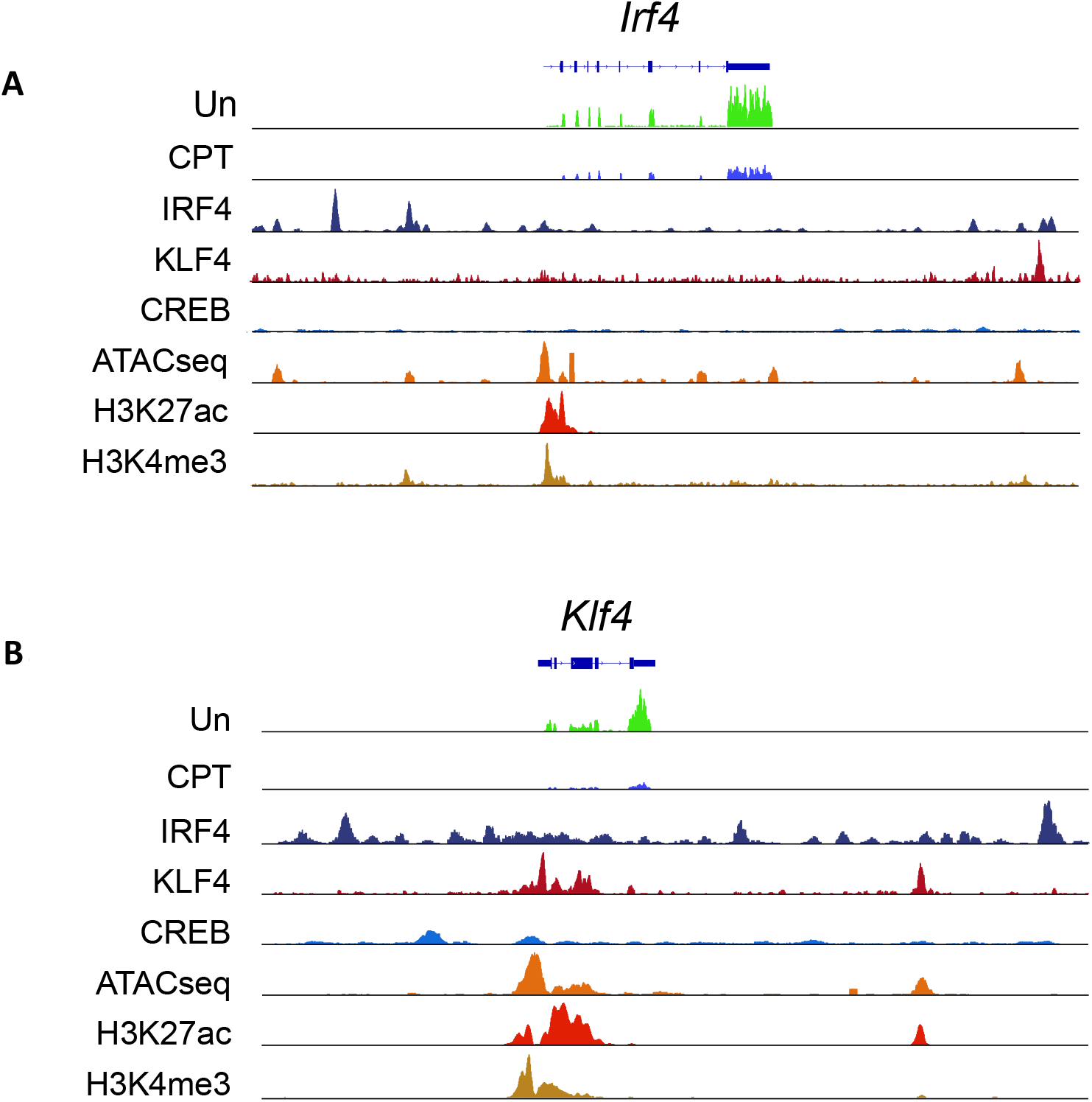
Expression from, chromatin structure of, and histone modifications at the *Irf4* and *Klf4* loci in cDC2s. **(A, B)** IRF4, KLF4 and CREB binding to the *Irf4* and *Klf4* loci and co-localization with open chromatin (ATACseq) and H3K27ac (enhancers and promoters) and H3K4me3 (promoters) modifications. The gene structures are shown at the top and RNAseq reads from untreated (Un) and CPT-treated (CPT) cDC2s are shown in green and blue.

**Supplemental Figure 10.**
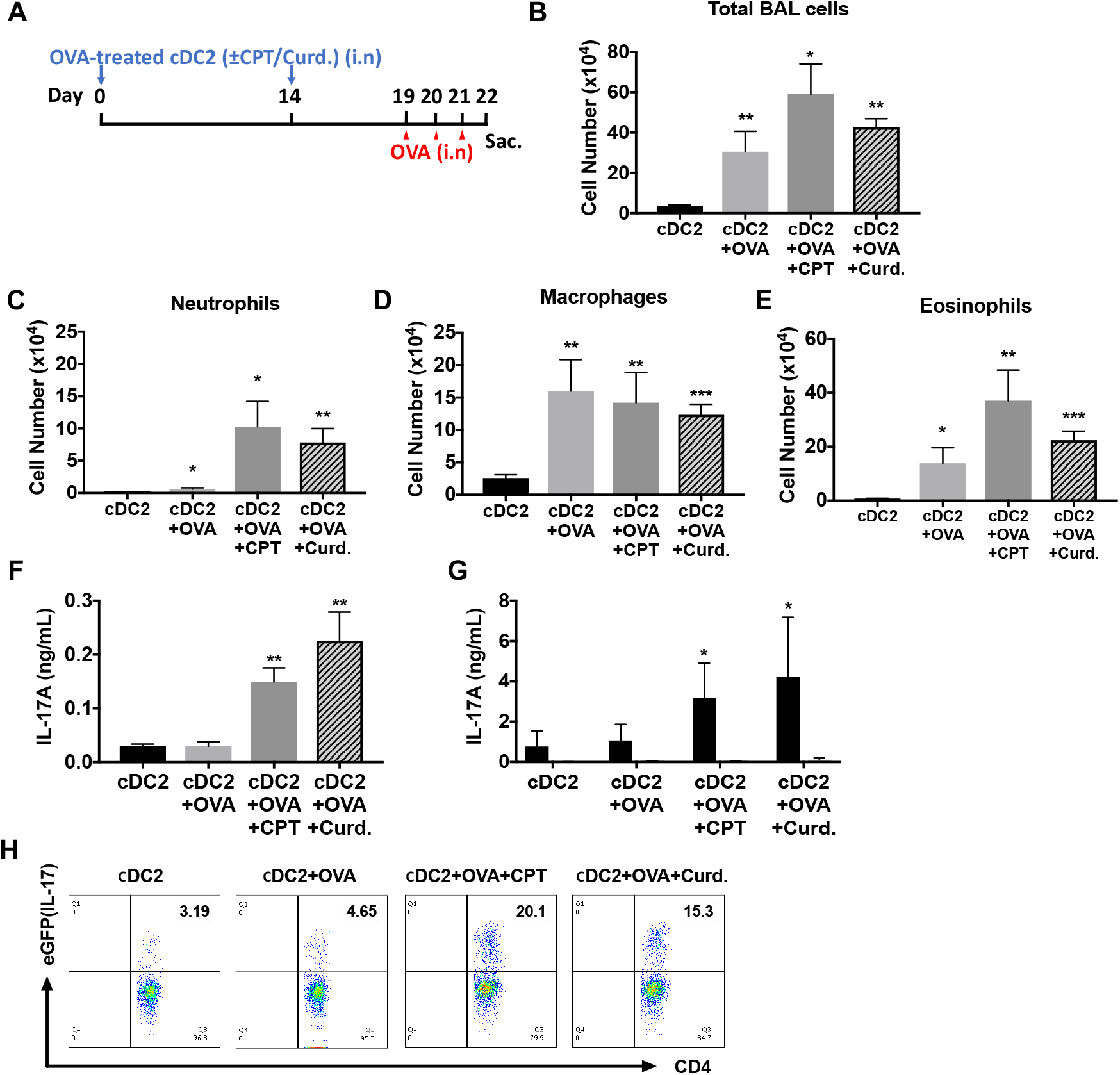
Adoptive transfer of OVA-pulsed, CPT or curdlan-treated cDC2s induces a Th17 bias and neutrophilic asthma in WT recipients. **(A)** Schematic of adoptive transfer protocol. WT cDC2s were incubated with OVA (200 μg/ml) in the presence of CPT or curdlan prior to i.n. transfer to WT (B6 mice) or IL-17-GFP mice (1×10^6^ cells/recipient) on day 0 and 14. OVA (50 µg/mouse) was used for the i.n. challenge on days 19, 20 and 21. **(B)** Total cell number, **(C)** neutrophils, **(D)** macrophages and **(E)** eosinophils number of BAL fluid. IL-17A levels from the **(F)** BAL fluid and **(G)** OVA (200 μg/ml)-stimulated lung cells. **(H)** Isolated lung cells from IL-17-GFP mice were analyzed (FACS) for the eGFP fluorescence intensity after PMA/ionomycin stimulation. Data are mean ± s.e.m, *n*=5 recipient mice in each group; * *p*<0.05, ** *p*<0.01.

## References

Agrawal, S., Gupta, S., & Agrawal, A. (2010). Human dendritic cells activated via dectin-1 are efficient at priming Th17, cytotoxic CD8 T and B cell responses. PLoS One, 5(10), e13418. doi:10.1371/journal.pone.0013418

Akira, S., Uematsu, S., & Takeuchi, O. (2006). Pathogen recognition and innate immunity. Cell, 124(4), 783–801. doi:10.1016/j.cell.2006.02.015

Anders, S., & Huber, W. (2010). Differential expression analysis for sequence count data. Genome Biol, 11(10), R106. doi:10.1186/gb-2010-11-10-r106

Bajana, S., Roach, K., Turner, S., Paul, J., & Kovats, S. (2012). IRF4 promotes cutaneous dendritic cell migration to lymph nodes during homeostasis and inflammation. J Immunol, 189(7), 3368–3377. doi:10.4049/jimmunol.1102613

Baranov, M., Ter Beest, M., Reinieren-Beeren, I., Cambi, A., Figdor, C. G., & van den Bogaart, G. (2014). Podosomes of dendritic cells facilitate antigen sampling. J Cell Sci, 127(Pt 5), 1052–1064. doi:10.1242/jcs.141226

Bat-Erdene, A., Miki, H., Oda, A., Nakamura, S., Teramachi, J., Amachi, R., … Abe, M. (2016). Synergistic targeting of Sp1, a critical transcription factor for myeloma cell growth and survival, by panobinostat and proteasome inhibitors. Oncotarget, 7(48), 79064–79075. doi:10.18632/oncotarget.12594

Bedoui, S., & Heath, W. R. (2015). Kruppel-ling of IRF4-Dependent DCs into Two Functionally Distinct DC Subsets. Immunity, 42(5), 785–787. doi:10.1016/j.immuni.2015.04.020

Berges, C., Naujokat, C., Tinapp, S., Wieczorek, H., Hoh, A., Sadeghi, M., … Daniel, V. (2005). A cell line model for the differentiation of human dendritic cells. Biochem Biophys Res Commun, 333(3), 896–907. doi:10.1016/j.bbrc.2005.05.171

Bharati, K., & Ganguly, N. K. (2011). Cholera toxin: a paradigm of a multifunctional protein. Indian J Med Res, 133, 179–187.

Biswas, P. S., Gupta, S., Stirzaker, R. A., Kumar, V., Jessberger, R., Lu, T. T., … Pernis, A. B. (2012). Dual regulation of IRF4 function in T and B cells is required for the coordination of T-B cell interactions and the prevention of autoimmunity. J Exp Med, 209(3), 581–596. doi:10.1084/jem.20111195

Boddicker, R. L., Kip, N. S., Xing, X., Zeng, Y., Yang, Z. Z., Lee, J. H., … Feldman, A. L. (2015). The oncogenic transcription factor IRF4 is regulated by a novel CD30/NF-kappaB positive feedback loop in peripheral T-cell lymphoma. Blood, 125(20), 3118–3127. doi:10.1182/blood-2014-05-578575

Bohme, J., Rossnagel, C., Jacobs, T., Behrends, J., Holscher, C., & Erdmann, H. (2016). Epstein-Barr virus-induced gene 3 suppresses T helper type 1, type 17 and type 2 immune responses after Trypanosoma cruzi infection and inhibits parasite replication by interfering with alternative macrophage activation. Immunology, 147(3), 338–348. doi:10.1111/imm.12565

Bovolenta, C., Driggers, P. H., Marks, M. S., Medin, J. A., Politis, A. D., Vogel, S. N., …, et al. (1994). Molecular interactions between interferon consensus sequence binding protein and members of the interferon regulatory factor family. Proc Natl Acad Sci U S A, 91(11), 5046–5050.

Brass, A. L., Kehrli, E., Eisenbeis, C. F., Storb, U., & Singh, H. (1996). Pip, a lymphoid-restricted IRF, contains a regulatory domain that is important for autoinhibition and ternary complex formation with the Ets factor PU.1. Genes Dev, 10(18), 2335–2347.

Cattaneo, F., Parisi, M., & Ammendola, R. (2013). Distinct signaling cascades elicited by different formyl peptide receptor 2 (FPR2) agonists. Int J Mol Sci, 14(4), 7193–7230. doi:10.3390/ijms14047193

Chen, Z., & O’Shea, J. J. (2008). Th17 cells: a new fate for differentiating helper T cells. Immunol Res, 41(2), 87–102. doi:10.1007/s12026-007-8014-9

Chung, Y., Yamazaki, T., Kim, B. S., Zhang, Y., Reynolds, J. M., Martinez, G. J., … Dong, C. (2013). Epstein Barr virus-induced 3 (EBI3) together with IL-12 negatively regulates T helper 17-mediated immunity to Listeria monocytogenes infection. PLoS Pathog, 9(9), e1003628. doi:10.1371/journal.ppat.1003628

Datta, S. K., Sabet, M., Nguyen, K. P., Valdez, P. A., Gonzalez-Navajas, J. M., Islam, S., Raz, E. (2010). Mucosal adjuvant activity of cholera toxin requires Th17 cells and protects against inhalation anthrax. Proc Natl Acad Sci U S A, 107(23), 10638–10643. doi:10.1073/pnas.1002348107

De Silva, N. S., Simonetti, G., Heise, N., & Klein, U. (2012). The diverse roles of IRF4 in late germinal center B-cell differentiation. Immunol Rev, 247(1), 73–92. doi:10.1111/j.1600-065X.2012.01113.x

Dobin, A., Davis, C. A., Schlesinger, F., Drenkow, J., Zaleski, C., Jha, S., … Gingeras, T. R. (2013). STAR: ultrafast universal RNA-seq aligner. Bioinformatics, 29(1), 15–21. doi:10.1093/bioinformatics/bts635

Doherty, T. A., Soroosh, P., Khorram, N., Fukuyama, S., Rosenthal, P., Cho, J. Y., … Croft, M. (2011). The tumor necrosis factor family member LIGHT is a target for asthmatic airway remodeling. Nat Med, 17(5), 596–603. doi:10.1038/nm.2356

Elcombe, S. E., Naqvi, S., Van Den Bosch, M. W., MacKenzie, K. F., Cianfanelli, F., Brown, G. D., & Arthur, J. S. (2013). Dectin-1 regulates IL-10 production via a MSK1/2 and CREB dependent pathway and promotes the induction of regulatory macrophage markers. PLoS One, 8(3), e60086. doi:10.1371/journal.pone.0060086

Gao, Y., Nish, S. A., Jiang, R., Hou, L., Licona-Limon, P., Weinstein, J. S., … Medzhitov, R. (2013). Control of T helper 2 responses by transcription factor IRF4-dependent dendritic cells. Immunity, 39(4), 722–732. doi:10.1016/j.immuni.2013.08.028

Gawden-Bone, C., Zhou, Z., King, E., Prescott, A., Watts, C., & Lucocq, J. (2010). Dendritic cell podosomes are protrusive and invade the extracellular matrix using metalloproteinase MMP-14. J Cell Sci, 123(Pt 9), 1427–1437. doi:10.1242/jcs.056515

Gerdes, N., Zhu, L., Ersoy, M., Hermansson, A., Hjemdahl, P., Hu, H., … Li, N. (2011). Platelets regulate CD4(+) T-cell differentiation via multiple chemokines in humans. Thromb Haemost, 106(2), 353–362. doi:10.1160/TH11-01-0020

Glass, D. B., Cheng, H. C., Mende-Mueller, L., Reed, J., & Walsh, D. A. (1989). Primary structural determinants essential for potent inhibition of cAMP-dependent protein kinase by inhibitory peptides corresponding to the active portion of the heat-stable inhibitor protein. J Biol Chem, 264(15), 8802–8810.

Gross, O., Gewies, A., Finger, K., Schafer, M., Sparwasser, T., Peschel, C., … Ruland, J. (2006). Card9 controls a non-TLR signalling pathway for innate anti-fungal immunity. Nature, 442(7103), 651–656. doi:10.1038/nature04926

Guilliams, M., Ginhoux, F., Jakubzick, C., Naik, S. H., Onai, N., Schraml, B. U., … Yona, S. (2014). Dendritic cells, monocytes and macrophages: a unified nomenclature based on ontogeny. Nat Rev Immunol, 14(8), 571–578. doi:10.1038/nri3712

Hambleton, S., Salem, S., Bustamante, J., Bigley, V., Boisson-Dupuis, S., Azevedo, J., … Gros, P. (2011). IRF8 mutations and human dendritic-cell immunodeficiency. N Engl J Med, 365(2), 127–138. doi:10.1056/NEJMoa1100066

Hansen, J. D., Vojtech, L. N., & Laing, K. J. (2011). Sensing disease and danger: a survey of vertebrate PRRs and their origins. Dev Comp Immunol, 35(9), 886–897. doi:10.1016/j.dci.2011.01.008

Hau, C. S., Kanda, N., Tada, Y., Shibata, S., Uozaki, H., Fukusato, T., … Watanabe, S. (2016). Lipocalin-2 exacerbates psoriasiform skin inflammation by augmenting T-helper 17 response. J Dermatol, 43(7), 785–794. doi:10.1111/1346-8138.13227

Hey, Y. Y., & O’Neill, H. C. (2012). Murine spleen contains a diversity of myeloid and dendritic cells distinct in antigen presenting function. J Cell Mol Med, 16(11), 2611–2619. doi:10.1111/j.1582-4934.2012.01608.x

Hirota, K., Duarte, J. H., Veldhoen, M., Hornsby, E., Li, Y., Cua, D. J., … Stockinger, B. (2011). Fate mapping of IL-17-producing T cells in inflammatory responses. Nat Immunol, 12(3), 255–263. doi:10.1038/ni.1993

Huber, M., & Lohoff, M. (2014). IRF4 at the crossroads of effector T-cell fate decision. Eur J Immunol, 44(7), 1886–1895. doi:10.1002/eji.201344279

Idzko, M., Ferrari, D., & Eltzschig, H. K. (2014). Nucleotide signalling during inflammation. Nature, 509(7500), 310–317. doi:10.1038/nature13085

Ikeda, S., Saijo, S., Murayama, M. A., Shimizu, K., Akitsu, A., & Iwakura, Y. (2014). Excess IL-1 signaling enhances the development of Th17 cells by downregulating TGF-beta-induced Foxp3 expression. J Immunol, 192(4), 1449–1458. doi:10.4049/jimmunol.1300387

Iwata, A., Durai, V., Tussiwand, R., Briseno, C. G., Wu, X., Grajales-Reyes, G. E., … Murphy, K. M. (2017). Quality of TCR signaling determined by differential affinities of enhancers for the composite BATF-IRF4 transcription factor complex. Nat Immunol, 18(5), 563–572. doi:10.1038/ni.3714

Jenuwein, T., & Allis, C. D. (2001). Translating the histone code. Science, 293(5532), 1074–1080. doi:10.1126/science.1063127

Kim, H. S., Park, K. H., Lee, H. K., Kim, J. S., Kim, Y. G., Lee, J. H., … Han, S. B. (2016). Curdlan activates dendritic cells through dectin-1 and toll-like receptor 4 signaling. Int Immunopharmacol, 39, 71–78. doi:10.1016/j.intimp.2016.07.013

Klein, U., Casola, S., Cattoretti, G., Shen, Q., Lia, M., Mo, T., … Dalla-Favera, R. (2006). Transcription factor IRF4 controls plasma cell differentiation and class-switch recombination. Nat Immunol, 7(7), 773–782. doi:10.1038/ni1357

Koria, P., Bhushan, A., Irimia, D., & Yarmush, M. L. (2012). Microfluidic Device for Examining Directional Sensing in Dendritic Cell Chemotaxis. Nano Life, 2(2). doi:10.1142/S1793984411000475

Koski, G. K., Schwartz, G. N., Weng, D. E., Gress, R. E., Engels, F. H., Tsokos, M., … Cohen, P. A. (1999). Calcium ionophore-treated myeloid cells acquire many dendritic cell characteristics independent of prior differentiation state, transformation status, or sensitivity to biologic agents. Blood, 94(4), 1359–1371.

Krausgruber, T., Blazek, K., Smallie, T., Alzabin, S., Lockstone, H., Sahgal, N., … Udalova, I. A. (2011). IRF5 promotes inflammatory macrophage polarization and TH1-TH17 responses. Nat Immunol, 12(3), 231–238. doi:10.1038/ni.1990

Krishnamoorthy, V., Kannanganat, S., Maienschein-Cline, M., Cook, S. L., Chen, J., Bahroos, N., … Sciammas, R. (2017). The IRF4 Gene Regulatory Module Functions as a Read-Write Integrator to Dynamically Coordinate T Helper Cell Fate. Immunity, 47(3), 481–497 e487. doi:10.1016/j.immuni.2017.09.001

Kumar, P., Chen, K., & Kolls, J. K. (2013). Th17 cell based vaccines in mucosal immunity. Curr Opin Immunol, 25(3), 373–380. doi:10.1016/j.coi.2013.03.011

Kurakula, K., Vos, M., Logiantara, A., Roelofs, J. J., Nieuwenhuis, M. A., Koppelman, G. H., … de Vries, C. J. (2015). Nuclear Receptor Nur77 Attenuates Airway Inflammation in Mice by Suppressing NF-kappaB Activity in Lung Epithelial Cells. J Immunol, 195(4), 1388–1398. doi:10.4049/jimmunol.1401714

Lambrecht, B. N., & Hammad, H. (2012). Lung dendritic cells in respiratory viral infection and asthma: from protection to immunopathology. Annu Rev Immunol, 30, 243–270. doi:10.1146/annurev-immunol-020711-075021

Lambrecht, B. N., Pauwels, R. A., & Fazekas De St Groth, B. (2000). Induction of rapid T cell activation, division, and recirculation by intratracheal injection of dendritic cells in a TCR transgenic model. J Immunol, 164(6), 2937–2946.

Leceta, J., Gomariz, R. P., Martinez, C., Abad, C., Ganea, D., & Delgado, M. (2000). Receptors and transcriptional factors involved in the anti-inflammatory activity of VIP and PACAP. Ann N Y Acad Sci, 921, 92–102.

Lee, J., Kim, T. H., Murray, F., Li, X., Choi, S. S., Broide, D. H., … Raz, E. (2015). Cyclic AMP concentrations in dendritic cells induce and regulate Th2 immunity and allergic asthma. Proc Natl Acad Sci U S A, 112(5), 1529–1534. doi:10.1073/pnas.1417972112

Lehtonen, A., Veckman, V., Nikula, T., Lahesmaa, R., Kinnunen, L., Matikainen, S., & Julkunen, I. (2005). Differential expression of IFN regulatory factor 4 gene in human monocyte-derived dendritic cells and macrophages. J Immunol, 175(10), 6570–6579.

LeibundGut-Landmann, S., Gross, O., Robinson, M. J., Osorio, F., Slack, E. C., Tsoni, S. V., … Reis e Sousa, C. (2007). Syk- and CARD9-dependent coupling of innate immunity to the induction of T helper cells that produce interleukin 17. Nat Immunol, 8(6), 630–638. doi:10.1038/ni1460

Liu, L., Yen, J. H., & Ganea, D. (2007). A novel VIP signaling pathway in T cells cAMP-->protein tyrosine phosphatase (SHP-2?)-->JAK2/STAT4-->Th1 differentiation. Peptides, 28(9), 1814–1824. doi:10.1016/j.peptides.2007.03.015

Lohning, M., Stroehmann, A., Coyle, A. J., Grogan, J. L., Lin, S., Gutierrez-Ramos, J. C., … Kamradt, T. (1998). T1/ST2 is preferentially expressed on murine Th2 cells, independent of interleukin 4, interleukin 5, and interleukin 10, and important for Th2 effector function. Proc Natl Acad Sci U S A, 95(12), 6930–6935.

Machida, I., Matsuse, H., Kondo, Y., Kawano, T., Saeki, S., Tomari, S., … Kohno, S. (2004). Cysteinyl leukotrienes regulate dendritic cell functions in a murine model of asthma. J Immunol, 172(3), 1833–1838.

Man, K., Miasari, M., Shi, W., Xin, A., Henstridge, D. C., Preston, S., … Kallies, A. (2013). The transcription factor IRF4 is essential for TCR affinity-mediated metabolic programming and clonal expansion of T cells. Nat Immunol, 14(11), 1155–1165. doi:10.1038/ni.2710

Marinissen, M. J., & Gutkind, J. S. (2001). G-protein-coupled receptors and signaling networks: emerging paradigms. Trends Pharmacol Sci, 22(7), 368–376.

Matsuyama, T., Grossman, A., Mittrucker, H. W., Siderovski, D. P., Kiefer, F., Kawakami, T., … Mak, T. W. (1995). Molecular cloning of LSIRF, a lymphoid-specific member of the interferon regulatory factor family that binds the interferon-stimulated response element (ISRE). Nucleic Acids Res, 23(12), 2127–2136.

Meinken, J., Walker, G., Cooper, C. R., & Min, X. J. (2015). MetazSecKB: the human and animal secretome and subcellular proteome knowledgebase. Database (Oxford), 2015. doi:10.1093/database/bav077

Merad, M., Sathe, P., Helft, J., Miller, J., & Mortha, A. (2013). The dendritic cell lineage: ontogeny and function of dendritic cells and their subsets in the steady state and the inflamed setting. Annu Rev Immunol, 31, 563–604. doi:10.1146/annurev-immunol-020711-074950

Mildner, A., & Jung, S. (2014). Development and function of dendritic cell subsets. Immunity, 40(5), 642–656. doi:10.1016/j.immuni.2014.04.016

Mitra, B., Jindal, R., Lee, S., Xu Dong, D., Li, L., Sharma, N., … Yarmush, M. L. (2013). Microdevice integrating innate and adaptive immune responses associated with antigen presentation by dendritic cells. RSC Adv, 3(36), 16002–16010. doi:10.1039/C3RA41308J

Mittrucker, H. W., Matsuyama, T., Grossman, A., Kundig, T. M., Potter, J., Shahinian, A., … Mak, T. W. (1997). Requirement for the transcription factor LSIRF/IRF4 for mature B and T lymphocyte function. Science, 275(5299), 540–543.

Mohrs, M., Shinkai, K., Mohrs, K., & Locksley, R. M. (2001). Analysis of type 2 immunity in vivo with a bicistronic IL-4 reporter. Immunity, 15(2), 303–311.

Mori Sequeiros Garcia, M., Gorostizaga, A., Brion, L., Gonzalez-Calvar, S. I., & Paz, C. (2015). cAMP-activated Nr4a1 expression requires ERK activity and is modulated by MAPK phosphatase-1 in MA-10 Leydig cells. Mol Cell Endocrinol, 408, 45–52. doi:10.1016/j.mce.2015.01.041

Muppala, S., Frolova, E., Xiao, R., Krukovets, I., Yoon, S., Hoppe, G., … Stenina-Adognravi, O. (2015). Proangiogenic Properties of Thrombospondin-4. Arterioscler Thromb Vasc Biol, 35(9), 1975–1986. doi:10.1161/ATVBAHA.115.305912

Murphy, K. M. (2013). Transcriptional control of dendritic cell development. Adv Immunol, 120, 239–267. doi:10.1016/B978-0-12-417028-5.00009-0

Myers, D. R., Lau, T., Markegard, E., Lim, H. W., Kasler, H., Zhu, M., … Roose, J. P. (2017). Tonic LAT-HDAC7 Signals Sustain Nur77 and Irf4 Expression to Tune Naive CD4 T Cells. Cell Rep, 19(8), 1558–1571. doi:10.1016/j.celrep.2017.04.076

Nagaoka, M., Yashiro, T., Uchida, Y., Ando, T., Hara, M., Arai, H., … Nishiyama, C. (2017). The Orphan Nuclear Receptor NR4A3 Is Involved in the Function of Dendritic Cells. J Immunol, 199(8), 2958–2967. doi:10.4049/jimmunol.1601911

Nakae, S., Komiyama, Y., Nambu, A., Sudo, K., Iwase, M., Homma, I., … Iwakura, Y. (2002). Antigen-specific T cell sensitization is impaired in IL-17-deficient mice, causing suppression of allergic cellular and humoral responses. Immunity, 17(3), 375–387.

Nakayama, T., Hirahara, K., Onodera, A., Endo, Y., Hosokawa, H., Shinoda, K., … Okamoto, Y. (2017). Th2 Cells in Health and Disease. Annu Rev Immunol, 35, 53–84. doi:10.1146/annurev-immunol-051116-052350

Nayar, R., Schutten, E., Bautista, B., Daniels, K., Prince, A. L., Enos, M., … Berg, L. J. (2014). Graded levels of IRF4 regulate CD8+ T cell differentiation and expansion, but not attrition, in response to acute virus infection. J Immunol, 192(12), 5881–5893. doi:10.4049/jimmunol.1303187

Nowyhed, H. N., Huynh, T. R., Thomas, G. D., Blatchley, A., & Hedrick, C. C. (2015). Cutting Edge: The Orphan Nuclear Receptor Nr4a1 Regulates CD8+ T Cell Expansion and Effector Function through Direct Repression of Irf4. J Immunol, 195(8), 3515–3519. doi:10.4049/jimmunol.1403027

Ochiai, K., Maienschein-Cline, M., Simonetti, G., Chen, J., Rosenthal, R., Brink, R., … Sciammas, R. (2013). Transcriptional regulation of germinal center B and plasma cell fates by dynamical control of IRF4. Immunity, 38(5), 918–929. doi:10.1016/j.immuni.2013.04.009

Palm, N. W., & Medzhitov, R. (2009). Pattern recognition receptors and control of adaptive immunity. Immunol Rev, 227(1), 221–233. doi:10.1111/j.1600-065X.2008.00731.x

Panasevich, S., Melen, E., Hallberg, J., Bergstrom, A., Svartengren, M., Pershagen, G., & Nyberg, F. (2013). Investigation of novel genes for lung function in children and their interaction with tobacco smoke exposure: a preliminary report. Acta Paediatr, 102(5), 498–503. doi:10.1111/apa.12204

Parnell, E., Palmer, T. M., & Yarwood, S. J. (2015). The future of EPAC-targeted therapies: agonism versus antagonism. Trends Pharmacol Sci, 36(4), 203–214. doi:10.1016/j.tips.2015.02.003

Pearce, E. J., & Everts, B. (2015). Dendritic cell metabolism. Nat Rev Immunol, 15(1), 18–29. doi:10.1038/nri3771

Persson, E. K., Uronen-Hansson, H., Semmrich, M., Rivollier, A., Hagerbrand, K., Marsal, J., … Agace, W. W. (2013). IRF4 transcription-factor-dependent CD103(+)CD11b(+) dendritic cells drive mucosal T helper 17 cell differentiation. Immunity, 38(5), 958–969. doi:10.1016/j.immuni.2013.03.009

Robinson, J. T., Thorvaldsdottir, H., Winckler, W., Guttman, M., Lander, E. S., Getz, G., & Mesirov, J. P. (2011). Integrative genomics viewer. Nat Biotechnol, 29(1), 24–26. doi:10.1038/nbt.1754

Rogers, N. C., Slack, E. C., Edwards, A. D., Nolte, M. A., Schulz, O., Schweighoffer, E., … Reis e Sousa, C. (2005). Syk-dependent cytokine induction by Dectin-1 reveals a novel pattern recognition pathway for C type lectins. Immunity, 22(4), 507–517. doi:10.1016/j.immuni.2005.03.004

Rogier, R., Ederveen, T. H. A., Boekhorst, J., Wopereis, H., Scher, J. U., Manasson, J., … Abdollahi-Roodsaz, S. (2017). Aberrant intestinal microbiota due to IL-1 receptor antagonist deficiency promotes IL-17- and TLR4-dependent arthritis. Microbiome, 5(1), 63. doi:10.1186/s40168-017-0278-2

Satoh, T., Takeuchi, O., Vandenbon, A., Yasuda, K., Tanaka, Y., Kumagai, Y., … Akira, S. (2010). The Jmjd3-Irf4 axis regulates M2 macrophage polarization and host responses against helminth infection. Nat Immunol, 11(10), 936–944. doi:10.1038/ni.1920

Schlitzer, A., McGovern, N., Teo, P., Zelante, T., Atarashi, K., Low, D., … Ginhoux, F. (2013). IRF4 transcription factor-dependent CD11b+ dendritic cells in human and mouse control mucosal IL-17 cytokine responses. Immunity, 38(5), 970–983. doi:10.1016/j.immuni.2013.04.011

Sciammas, R., Shaffer, A. L., Schatz, J. H., Zhao, H., Staudt, L. M., & Singh, H. (2006). Graded expression of interferon regulatory factor-4 coordinates isotype switching with plasma cell differentiation. Immunity, 25(2), 225–236. doi:10.1016/j.immuni.2006.07.009

Shaffer, A. L., Emre, N. C., Lamy, L., Ngo, V. N., Wright, G., Xiao, W., Staudt, L. M. (2008). IRF4 addiction in multiple myeloma. Nature, 454(7201), 226–231. doi:10.1038/nature07064

Shaffer, A. L., Emre, N. C., Romesser, P. B., & Staudt, L. M. (2009). IRF4: Immunity. Malignancy! Therapy? Clin Cancer Res, 15(9), 2954–2961. doi:10.1158/1078-0432.CCR-08-1845

Sharma, S., Grandvaux, N., Mamane, Y., Genin, P., Azimi, N., Waldmann, T., & Hiscott, J. (2002). Regulation of IFN regulatory factor 4 expression in human T cell leukemia virus-I-transformed T cells. J Immunol, 169(6), 3120–3130.

Shukla, V., Ma, S., Hardy, R. R., Joshi, S. S., & Lu, R. (2013). A role for IRF4 in the development of CLL. Blood, 122(16), 2848–2855. doi:10.1182/blood-2013-03-492769

Sichien, D., Lambrecht, B. N., Guilliams, M., & Scott, C. L. (2017). Development of conventional dendritic cells: from common bone marrow progenitors to multiple subsets in peripheral tissues. Mucosal Immunol, 10(4), 831–844. doi:10.1038/mi.2017.8

Sriram, K., & Insel, P. A. (2018). G Protein-Coupled Receptors as Targets for Approved Drugs: How Many Targets and How Many Drugs? Mol Pharmacol, 93(4), 251–258. doi:10.1124/mol.117.111062

Suzuki, S., Honma, K., Matsuyama, T., Suzuki, K., Toriyama, K., Akitoyo, I., … Kumatori, A. (2004). Critical roles of interferon regulatory factor 4 in CD11bhighCD8alpha-dendritic cell development. Proc Natl Acad Sci U S A, 101(24), 8981–8986. doi:10.1073/pnas.0402139101

Takaoka, A., Yanai, H., Kondo, S., Duncan, G., Negishi, H., Mizutani, T., … Taniguchi, T. (2005). Integral role of IRF-5 in the gene induction programme activated by Toll-like receptors. Nature, 434(7030), 243–249. doi:10.1038/nature03308

Takeuchi, O., & Akira, S. (2010). Pattern recognition receptors and inflammation. Cell, 140(6), 805–820. doi:10.1016/j.cell.2010.01.022

Tezuka, T., Ogawa, H., Azuma, M., Goto, H., Uehara, H., Aono, Y., … Nishioka, Y. (2015). IMD-4690, a novel specific inhibitor for plasminogen activator inhibitor-1, reduces allergic airway remodeling in a mouse model of chronic asthma via regulating angiogenesis and remodeling-related mediators. PLoS One, 10(3), e0121615. doi:10.1371/journal.pone.0121615

Tussiwand, R., Everts, B., Grajales-Reyes, G. E., Kretzer, N. M., Iwata, A., Bagaitkar, J., … Murphy, K. M. (2015). Klf4 expression in conventional dendritic cells is required for T helper 2 cell responses. Immunity, 42(5), 916–928. doi:10.1016/j.immuni.2015.04.017

Urushima, H., Fujimoto, M., Mishima, T., Ohkawara, T., Honda, H., Lee, H., … Naka, T. (2017). Leucine-rich alpha 2 glycoprotein promotes Th17 differentiation and collagen-induced arthritis in mice through enhancement of TGF-beta-Smad2 signaling in naive helper T cells. Arthritis Res Ther, 19(1), 137. doi:10.1186/s13075-017-1349-2

Vander Lugt, B., Khan, A. A., Hackney, J. A., Agrawal, S., Lesch, J., Zhou, M., … Singh, H. (2014). Transcriptional programming of dendritic cells for enhanced MHC class II antigen presentation. Nat Immunol, 15(2), 161–167. doi:10.1038/ni.2795

Wang, S., Sun, H., Ma, J., Zang, C., Wang, C., Wang, J., … Liu, X. S. (2013). Target analysis by integration of transcriptome and ChIP-seq data with BETA. Nat Protoc, 8(12), 2502–2515. doi:10.1038/nprot.2013.150

Wang, X., Zhang, Y., Wang, Z., Liu, X., Zhu, G., Han, G., … Wang, R. (2018). AntiIL39 (IL23p19/Ebi3) polyclonal antibodies ameliorate autoimmune symptoms in lupuslike mice. Mol Med Rep, 17(1), 1660–1666. doi:10.3892/mmr.2017.8048

Wang, Y. H., Voo, K. S., Liu, B., Chen, C. Y., Uygungil, B., Spoede, W., … Liu, Y. J. (2010). A novel subset of CD4(+) T(H)2 memory/effector cells that produce inflammatory IL-17 cytokine and promote the exacerbation of chronic allergic asthma. J Exp Med, 207(11), 2479–2491. doi:10.1084/jem.20101376

Watkins, A. A., Yasuda, K., Wilson, G. E., Aprahamian, T., Xie, Y., Maganto-Garcia, E., … Rifkin, I. R. (2015). IRF5 deficiency ameliorates lupus but promotes atherosclerosis and metabolic dysfunction in a mouse model of lupus-associated atherosclerosis. J Immunol, 194(4), 1467–1479. doi:10.4049/jimmunol.1402807

Wehbi, V. L., & Tasken, K. (2016). Molecular Mechanisms for cAMP-Mediated Immunoregulation in T cells - Role of Anchored Protein Kinase A Signaling Units. Front Immunol, 7, 222. doi:10.3389/fimmu.2016.00222

Whyte, W. A., Orlando, D. A., Hnisz, D., Abraham, B. J., Lin, C. Y., Kagey, M. H., … Young, R. A. (2013). Master transcription factors and mediator establish super-enhancers at key cell identity genes. Cell, 153(2), 307–319. doi:10.1016/j.cell.2013.03.035

Williams, J. W., Tjota, M. Y., Clay, B. S., Vander Lugt, B., Bandukwala, H. S., Hrusch, C. L., … Sperling, A. I. (2013). Transcription factor IRF4 drives dendritic cells to promote Th2 differentiation. Nat Commun, 4, 2990. doi:10.1038/ncomms3990

Worbs, T., Hammerschmidt, S. I., & Forster, R. (2017). Dendritic cell migration in health and disease. Nat Rev Immunol, 17(1), 30–48. doi:10.1038/nri.2016.116

Xia, C. Q., & Kao, K. J. (2003). Effect of CXC chemokine platelet factor 4 on differentiation and function of monocyte-derived dendritic cells. Int Immunol, 15(8), 1007–1015.

Xie, F., Li, B. X., Kassenbrock, A., Xue, C., Wang, X., Qian, D. Z., … Xiao, X. (2015). Identification of a Potent Inhibitor of CREB-Mediated Gene Transcription with Efficacious in Vivo Anticancer Activity. J Med Chem, 58(12), 5075–5087. doi:10.1021/acs.jmedchem.5b00468

Yang, K., Li, Q., & Zhao, Y. N. (2007). [Research on A23187 inducing HL-60 cells to differentiate into dendritic cells]. Sichuan Da Xue Xue Bao Yi Xue Ban, 38(2), 209–212.

Yao, S., Buzo, B. F., Pham, D., Jiang, L., Taparowsky, E. J., Kaplan, M. H., & Sun, J. (2013). Interferon regulatory factor 4 sustains CD8(+) T cell expansion and effector differentiation. Immunity, 39(5), 833–845. doi:10.1016/j.immuni.2013.10.007

Yoshitomi, H., Sakaguchi, N., Kobayashi, K., Brown, G. D., Tagami, T., Sakihama, T., … Sakaguchi, S. (2005). A role for fungal {beta}-glucans and their receptor Dectin-1 in the induction of autoimmune arthritis in genetically susceptible mice. J Exp Med, 201(6), 949–960. doi:10.1084/jem.20041758

